# Comprehensive multimodal and multiomic profiling reveals epigenetic and transcriptional reprogramming in lung tumors

**DOI:** 10.1101/2024.06.06.597667

**Authors:** Peiyao Wu, Zhengzhi Liu, Lina Zheng, Zirui Zhou, Wei Wang, Chang Lu

## Abstract

Epigenomic mechanisms are critically involved in mediation of genetic and environmental factors that underlie cancer development. Histone modifications represent highly informative epigenomic marks that reveal activation and repression of gene activities and dysregulation of transcriptional control due to tumorigenesis. Here, we present a comprehensive epigenomic and transcriptomic mapping of 18 tumor and 20 non-neoplastic tissues from non-small cell lung adenocarcinoma patients. Our profiling covers 5 histone marks including activating (H3K4me3, H3K4me1, and H3K27ac) and repressive (H3K27me3 and H3K9me3) marks and the transcriptome using only 20 mg of tissue per sample, enabled by low-input omic technologies. Using advanced integrative bioinformatic analysis, we uncovered cancer-driving signaling cascade networks, changes in 3D genome modularity, and differential expression and functionalities of transcription factors and noncoding RNAs. Many of these identified genes and regulatory molecules showed no significant change in their expression or a single epigenomic modality, emphasizing the power of integrative multimodal and multiomic analysis using patient samples.

## Introduction

Although transcription factor (TF) networks generally define cellular states, chromatin landscape often determines the permissiveness of a switch between different cellular states ^1,2^. Early variations in the epigenetic landscape due to histone modifications, DNA methylation, and non-coding RNA bindings in a premalignant cell potentially set the stage for aberrant TF activities and signaling pathways associated with oncogenic transformation ^3,4^. Canonical chromatin states are marked by specific histone modifications (e.g. H3K4me3 and H3K27ac for active states; H3K9me3 and H3K27me3 for repressive states) ^3^. The integrative analysis of chromatin state, gene expression, and long-range interactions allows identification of driver TFs and construction of TF networks ^5^.

Non-small cell lung cancer (NSCLC) is the leading cause of cancer mortality and accounts for 85% of all lung cancers ^6^. Although there has been significant progress in diagnosis and therapy of NSCLC over the years, improved understanding of its molecular basis is still required for guiding therapeutic decisions. Profiling of primary tumor samples has revealed critical information on reprogramming of regulatory elements in cancer ^7,8^. Identifying epigenomic signatures is crucial for understanding NSCLC pathogenesis and selecting the optimal treatment strategy. Epigenetic-mediated tumor suppressor gene silencing via hypermethylation or histone code was implicated in lung cancer ^9^. Global level of specific histone marks has been linked with prognosis of lung cancers ^10,11^. In spite of the progress, genome-wide profiling of histone modification was rarely performed directly on human tumor tissues due to the low quantity of cells that can be isolated from these samples^12^. Xenograft and in vitro cultures had to be used when multimodal (>3) profiling was conducted ^7,13–16^. Despite insights gained from analyses of individual histone marks, a comprehensive characterization of the epigenomic landscape including activating and repressive marks in primary tumor cells is necessary to deepen the understanding of epigenomic dynamics involved in NSCLC pathogenesis.

In this work, we conducted comprehensive epigenomic and transcriptomic profiling of primary non-small cell lung tumors (N = 18, adenocarcinoma subtype) and non-neoplastic tissues (N= 20) using low-input technologies, MOWChIP-seq ^17–19^ and Smart-seq2 ^20^. Five key histone modification marks (active marks H3K4me3, H3K27ac, and H3K4me1; repressive marks H3K27me3 and H3K9me3), and gene expression were examined in replicates for each tissue sample. We identified numerous differentially modified epigenomic and transcriptomic regions between tumor and non-neoplastic tissues. Further analysis by integrating histone modification and transcription signal patterns identified 6 sections of genomic loci, with their signature cRAMs, which presented distinct functions in pathway analysis. Finally, important TFs in the cancer group and their regulatee genes were identified to function in the pathways associated with lung cancer development.

## Results

### Comprehensive multimodal and multiomic profiling using patient samples

We employed MOWChIP-seq^17,18^ to profile five histone modifications (H3K27ac, H3K4me3, H3K4me1, H3K27me3, and H3K9me3) and Smart-seq2 ^20^ for transcriptome profiling, with two isogenic replicates utilizing approximately 20 mg of tissue from each of 18 NSCLC samples (lung adenocarcinoma) and 20 non-neoplastic lung tissue samples (**Figure 1** and **Table S1**). 30,000 cells were used to create each ChIP-seq replicate, while 10,000 cells were used for each RNA-seq replicate.

**Figure 1.**
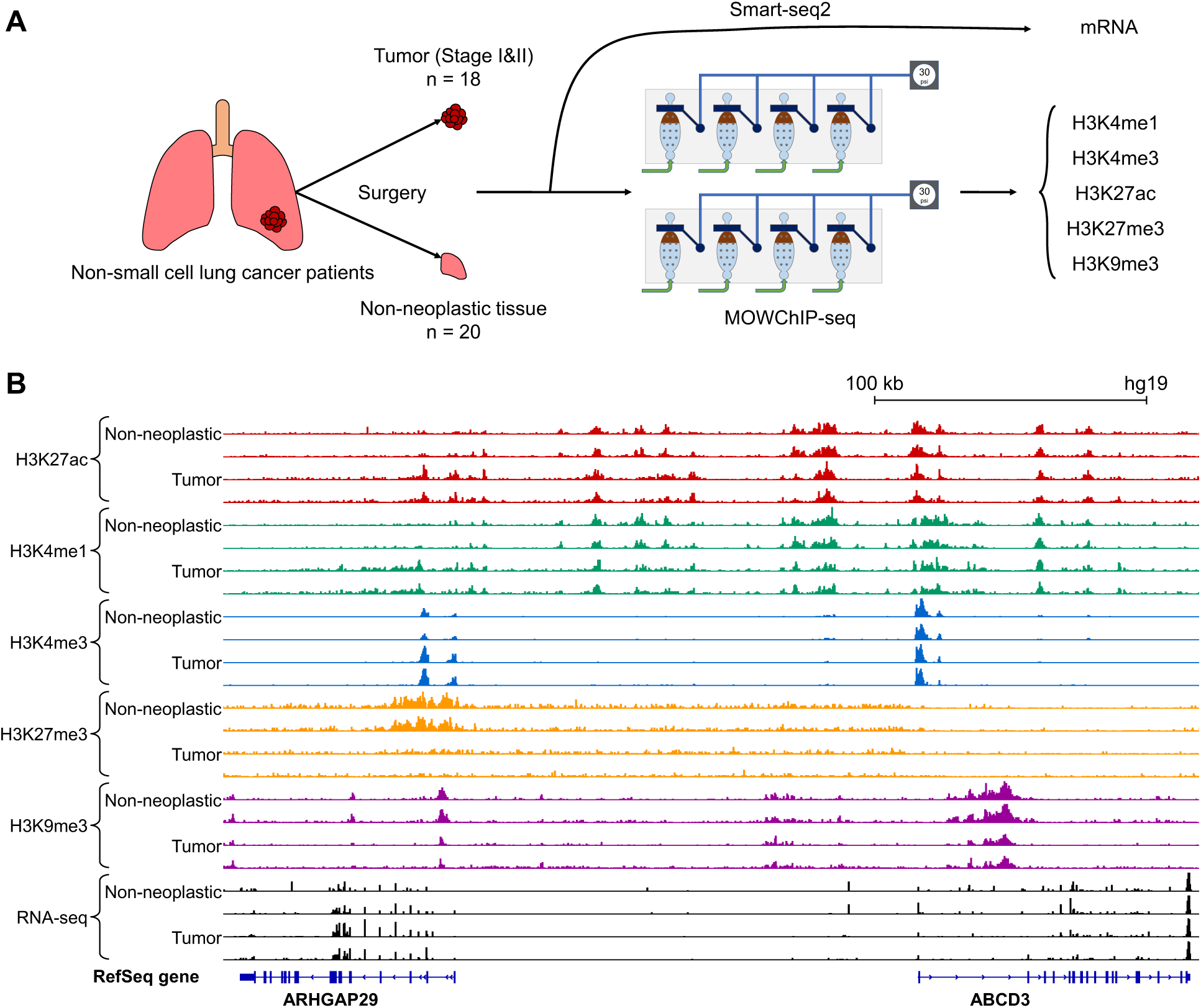
Overview of experiments. **(A)** Illustration of the experimental process. **(B)** Genome browser views of normalized ChIP-seq and RNA-seq signals from representative tumor and non-neoplastic samples.

Our MOWChIP-seq technology yielded average unique reads of approximately 12.2, 17.0, 12.2, 25.5, 37.8 million for histone modifications H3K27ac, H3K4me3, H3K4me1, H3K27me3, and H3K9me3, respectively (**Figure S1** and **Table S2**). Notably, MOWChIP-seq datasets showed minimal background noise ^18^, with the fraction of reads in called peaks (FRiP) averaging at 0.377, 0.450, 0.396, 0.122, and 0.181 for H3K27ac, H3K4me3, H3K4me1, H3K27me3, and H3K9me3, respectively (**Table S2**).

Furthermore, we calculated the normalized strand cross-correlation (NSC) and relative strand cross-correlation (RSC) to assess sequencing read enrichment around histone modification sites ^21^ (**Table S2**). The average NSC values were 1.12, 1.66, 1.13, 1.02, and 1.03, while the average RSC were 2.01, 1.60, 1.45, 3.87, and 1.47, for H3K27ac, H3K4me3, H3K4me1, H3K27me3, and H3K9me3, respectively. These NSC and RSC values exceeded or closely approached the recommended thresholds of 1.05 and 1.0, respectively, by ENCODE (https://genome.ucsc.edu/ENCODE/qualityMetrics.html). The Pearson’s correlations between replicates were consistently high, with averages of 0.93, 0.95, 0.93, 0.92, and 0.95 for H3K27ac, H3K4me3, H3K4me1, H3K27me3, and H3K9me3, respectively.

In our RNA-seq datasets, we obtained an average of approximately 11.1 million uniquely mapped reads, with an average mapping rate of 69.3% (**Figure S2** and **Table S2**). Additionally, the average GC content was 50.5%, and the exon percentage was 82.2%. All these results collectively demonstrated the successful generation of high-quality data through our low-input technologies.

### Differential analyses reveal widespread changes on activating and repressive histone marks and in transcriptome

To elucidate the epigenomic and transcriptomic changes associated with non-small cell lung cancer (NSCLC), we identified differentially modified epigenetic regions (DMERs) using DiffBind ^22^ and differentially expressed genes (DEGs) using DESeq2 ^23^ across the 18 tumor and 20 non-neoplastic lung tissue samples. Our analysis revealed significant alterations in histone modifications and gene expression profiles between the two groups.

We identified a total of 27,233 H3K27ac (13,415 gain and 13,818 loss of H3K27ac peaks in tumor compared to non-neoplastic samples), 44,677 H3K4me1 (21,132 gain and 23,545 loss), 13,005 H3K4me3 (8,725 gain and 4,280 loss), 5,091 H3K27me3 (2,877 gain and 2,214 loss), and 18,589 H3K9me3 (11,211 gain and 7,378 loss) regions that were differentially modified, along with 1,053 differentially expressed genes from RNA-seq (**Figure S3**).

The top ranked 96 differentially expressed genes (DEGs) identified from DESeq2 with FDR <= 0.0001 and |log2(fold change)| >= 1 showed distinct separation between tumor and non-neoplastic tissue groups (**Figure 2A**). These genes were significantly enriched in cancer and cell proliferation/apoptosis-related pathways (Benjamini-Hochberg FDR <= 0.05), as reported by the g:Profiler ^24^ web server (**Figure 2B**). We also investigated the functional pathways enriched in NSCLC-specific DMER landscapes (**Figure 2C**). Notably, general cancer-related terms such as “pathways in cancer” were recurrently observed in both DMERs and DEGs, indicating robust shifts in histone modification and gene expression profiles in NSCLC tumors. Several other high-ranking terms were shared among differential histone modifications and gene expression, including “Rap1 signaling pathway”, “Axon guidance”, “Focal adhesion”, “PI3K-Akt signaling pathway”, “Calcium signaling pathway”, “MAPK signaling pathway”, and “Ras signaling pathway”. These pathways play diverse yet interconnected roles in cancer progression. For instance, dysregulated expression of protein cues for axon guidance is implicated in lung cancer ^25^, while aberrations in regulatory genes of axon guidance are frequent in cancer ^26^. Calcium signaling pathway has been reported to be closely involved in the proliferation and apoptosis of NSCLC cells ^27^. In addition, pathways such as Rap1 and Ras signaling are particularly intriguing. Rap1 signaling plays diverse roles in tumor invasion and metastasis ^28^, and controls cell adhesion with its associated signaling network ^29^. PI3K-Akt signaling pathway is a downstream process of oncogenic Ras signaling, and both are targets for new therapies in NSCLC ^30,31^. MAPK signaling pathways are involved in tumorigenesis and related to Ras proteins through signaling networks ^32^. The robust detection of the linked pathways of Rap1, Ras, PI3K-Akt, MAPK, and focal adhesion suggests a network of signaling cascades affecting cell adhesion in NSCLC tumors. Moreover, focal adhesion has been shown to affect treatment resistance in cancer ^33^, and its associated signaling cascades may play crucial roles in the development and progression of NSCLC tumors.

**Figure 2.**
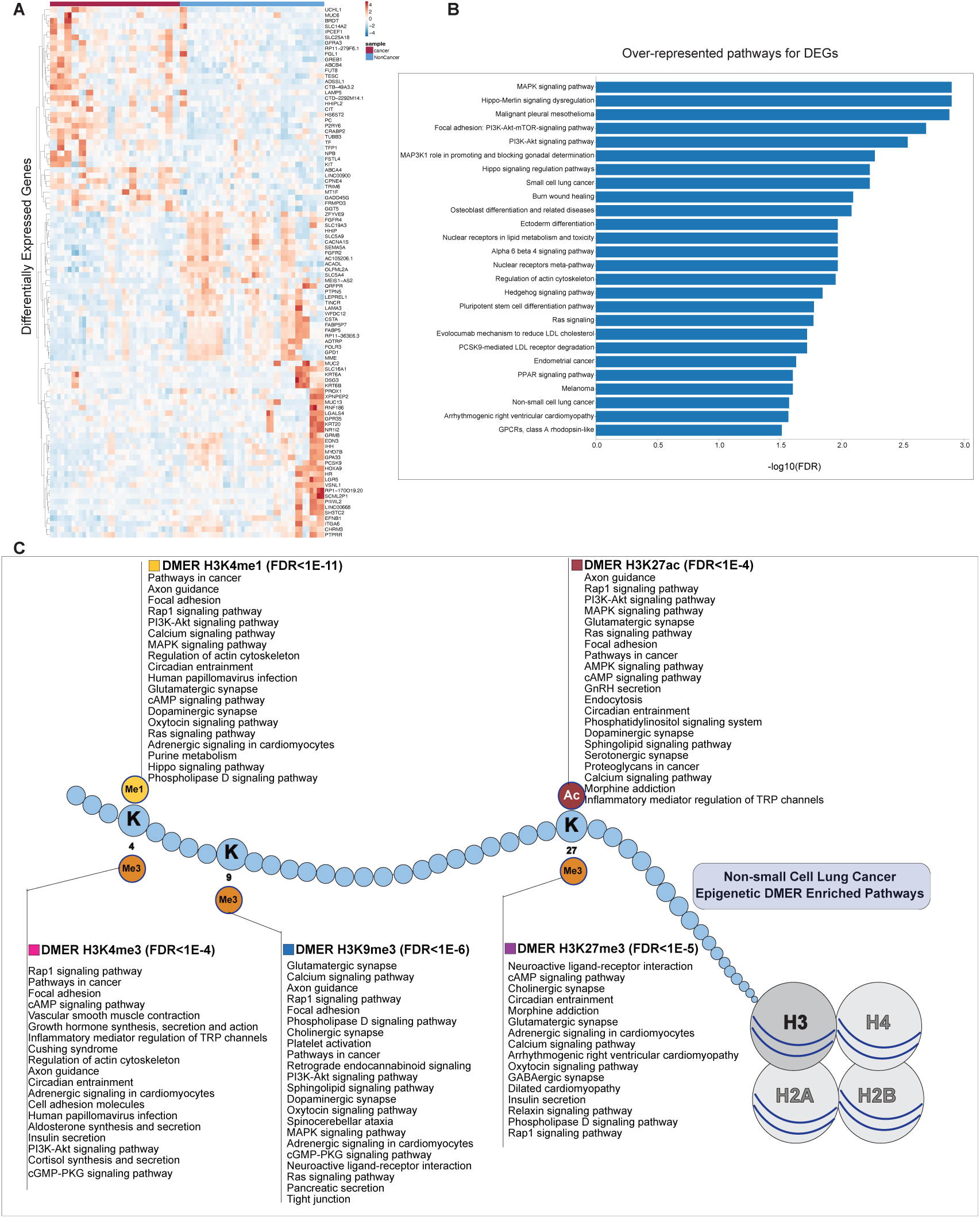
Differentially expressed genes and modified epigenetic regions. **(A)** Heatmap for the top-ranked differentially expressed genes (DEGs) between the tumor and non-neoplastic samples. **(B)** The over-represented pathways for the top ranked DEGs between tumor and non-neoplastic samples. **(C)** The enriched pathways for the differentially modified epigenetic regions (DMERs) for each individual histone mark (H3K27ac, H3K4me1, H3K4me3, H3K27me3, and H3K9me3).

### Clustering of genomic regions reveals the interdependent epigenomic and transcriptomic features due to NSCLC pathogenesis

To capture the comprehensive landscape of genomic regions co-modified across multiple histone marks and transcriptomic profiles for NSCLC and non-neoplastic lung tissues, we performed an unbiased unsupervised integrative analysis using EpiSig ^34,35^. EpiSig systematically scanned every 5kb-long region in the human genome to identify those with significant signals for any of the histone marks and gene expression across all samples, and subsequently clustered them based on similar epigenetic and transcriptomic patterns. Our EpiSig analysis generated 429 clusters, with an average number of 66 loci per cluster (**Figure 3**). We further categorized these clusters into six larger sections with distinct patterns through K-Means clustering (**Figure 3**).

**Figure 3.**
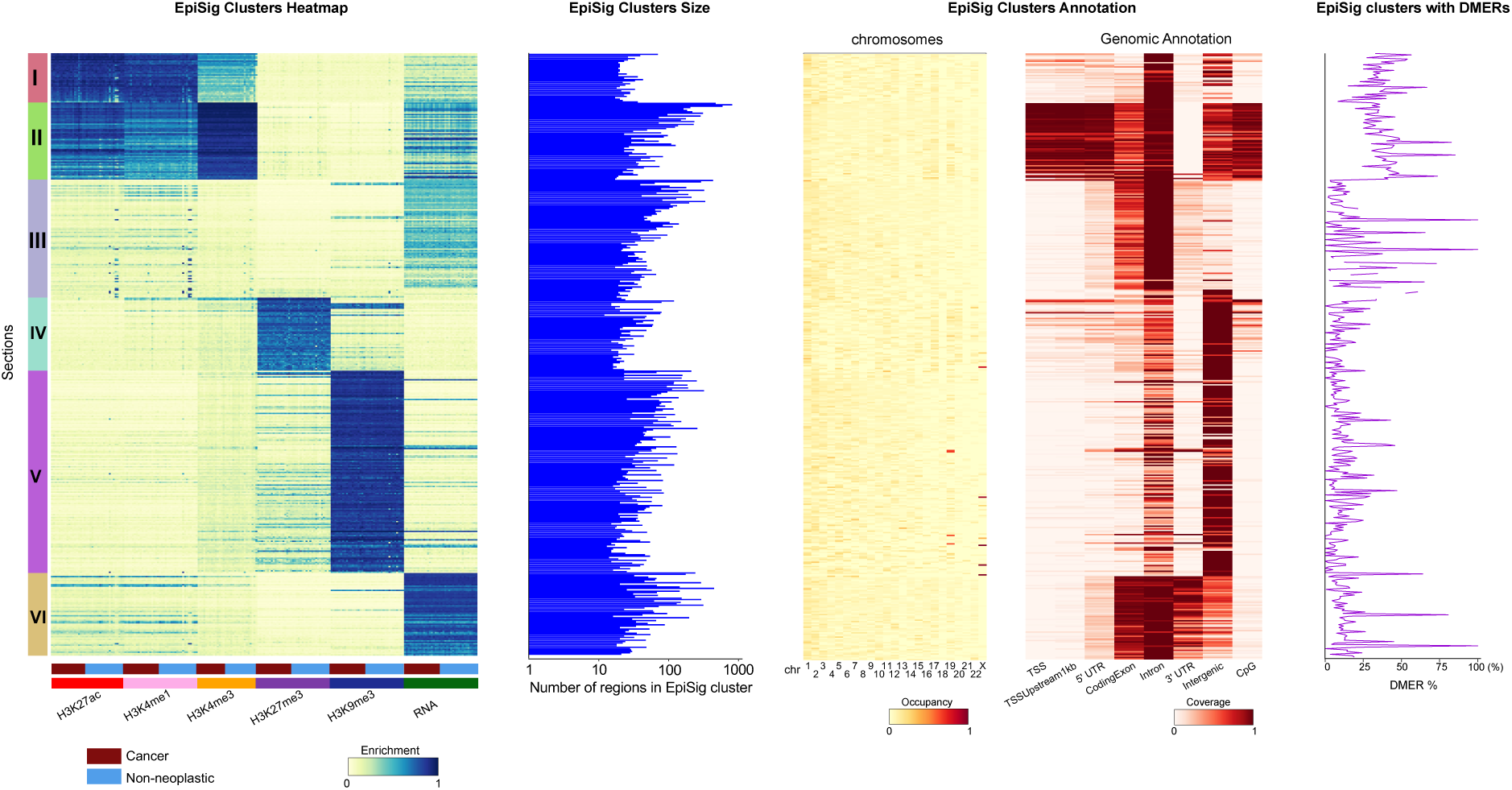
EpiSig integrative analysis on the multimodal epigenomic and transcriptomic signals. From left to right: Composition of EpiSig sections; Size distribution of EpiSig clusters; Genomic distribution of EpiSig clusters; Genomic annotation of EpiSig clusters; EpiSig clusters with DMERs.

Specifically, Section I comprised active enhancers, displaying high enrichment of H3K27ac and H3K4me1 and moderate levels of H3K4me3. Notably, Section I predominantly occupied intron and intergenic regions rather than promoters. In contrast, Section II represented active promoter regions, characterized by enrichment of H3K27ac and H3K4me3, along with moderate levels of H3K4me1 and RNA-seq signals, and largely occupied CpG islands, TSSs, TSS upstream 1 kb regions, 5’ UTRs, introns, and intergenic regions. Sections IV and V were identified as repressed regions. Section IV was marked primarily by H3K27me3 and weak H3K4me3, residing in the CpG islands and TSS regions; while Section V was enriched with the heterochromatin mark H3K9me3, accompanied by weak H3K27me3 and H3K4me3 signals, spanning intergenic regions. Notably, Section V was slightly biased towards chr19 and chrX, unlike the even chromosome distribution observed in other sections (**Figure 3**).

Sections III and VI represented transcribed regions, enriched with RNA-seq signals. Section III had slightly weaker RNA-seq but stronger H3K27ac/H3K4me1 signals than Section VI. Both sections encompassed gene body regions, including 5’ UTRs, coding exons, introns, and 3’UTRs. Additionally, sections I, II, III, and VI were highly enriched with DMERs/DEGs, compared to section IV and V (**Figure 3**). Specifically, Section I was enriched with H3K27ac and H3K4me1 DMERs, Section II with H3K27ac, H3K4me1, and H3K4me3 DMERs, Section III and Section VI with H3K27ac and H3K4me1 DMERs, Section IV with H3K27me3, H3K4me1, and RNA DMERs, and Section V with H3K9me3 and RNA DMERs (**Figure 3**)

This integrative analysis provides a comprehensive understanding of the genomic regions showing coordinated epigenetic and transcriptomic changes in NSCLC and non-neoplastic lung tissues, facilitating the identification of key regulatory elements involved in NSCLC pathogenesis.

### Integrative analysis reveals NSCLC-associated changes in chromatin organization

The alterations in global chromatin organization are recognized as key drivers of cancer progression ^36–38^. To explore the differences in chromatin organization between cancerous and non-neoplastic lung tissues, we employed findRAM method ^39^ to identify regulation associated modules (RAMs). RAMs represent genome-wide modular patterns indicative of chromatin activities, such as chromatin loops and super-enhancer clusters, at a better scale ^39^. Deletion of the RAM boundaries would severely perturb chromatin organization and affect cell fitness.

We used H3K27ac peak densities to identify RAMs in each individual sample using the sliding window strategy at a step size of 250 kb and a flanking window size of 500 kb (**Methods**), and 760 consensus RAMs (cRAMs) for the tumor group and 759 cRAMs for the non-neoplastic group were detected, respectively. **Figure 4A** displayed individual RAMs on chr6 for “13T” (tumor) and “13N” (non-neoplastic), as is shown, RAMs within the 65 - 75 Mb genomic region split in tumor samples. Among all the cRAMs, 682 were shared between both groups (**Figure 4B**). Similar to the previous reports ^39^, the median size of tumor cRAMs and non-neoplastic cRAMs were around 2.375 Mb and 2.25 Mb, respectively (**Figure S4)**. In addition to the shared cRAMs, there were 88 differential cRAM regions undergoing splitting or merging events, leading to 78 tumor-specific and 77 non-neoplastic-specific cRAMs, indicating the pathologically relevant chromatin organization changes (**Figure 4B**). To evaluate the functions of these differential cRAMs, we examined the differential binding signals (H3K27ac, H3K4me1, H3K4me3, H3K27me3, H3K9me3 and gene expression) within them. Interestingly, 87 out of 88 regions contained differential signals, with 97.8% of them displayed at least two differential signals (**Figure 4C**). **Figure 4D** showed an example region chr7:34,942,762-35,111,521 (hg19), where the differential cRAMs region was chr7:34,750,000-407,500,000. In this region, gene *DPY19L1* was observed to be differentially expressed and multiple H3K27ac, H3K4me3, and H3K4me1 differentially binding sites were discovered. **Figure 4E** displayed the pathways in the KEGG and Reactom databases that were enriched in the histone modification mark and gene expression signals in the differential cRAMs globally ^24,40,41^. Besides the common cancer-related pathways in the KEGG database as shown in **Figure 2C**, other pathways were also observed. For example, H3K27ac, H3K4me1, and H3K4me3 differential signals in the differential cRAMs were highly enriched in the ECM-receptor interaction pathway. ECM pathway and its remodeling by cancer have been substantiated to be highly associated with cancer cells growth, survival, and metastasis, and thus it would be a prognostic signature of non-small lung cancer ^42–45^. In addition, we observed pathways such as “TGFBR1KD mutants in cancer”, “loss of function of TGFBR1 in cancer”, “loss of function of smad2/3 in cancer”, and “signaling by TGF-beta receptor complex in cancer”, which are critical in cancer development and progress and considered therapeutic targets in clinical studies ^46,47^. Function analysis for EpiSig sections over the 88 differential cRAMs were displayed in **Figure S5**.

**Figure 4.**
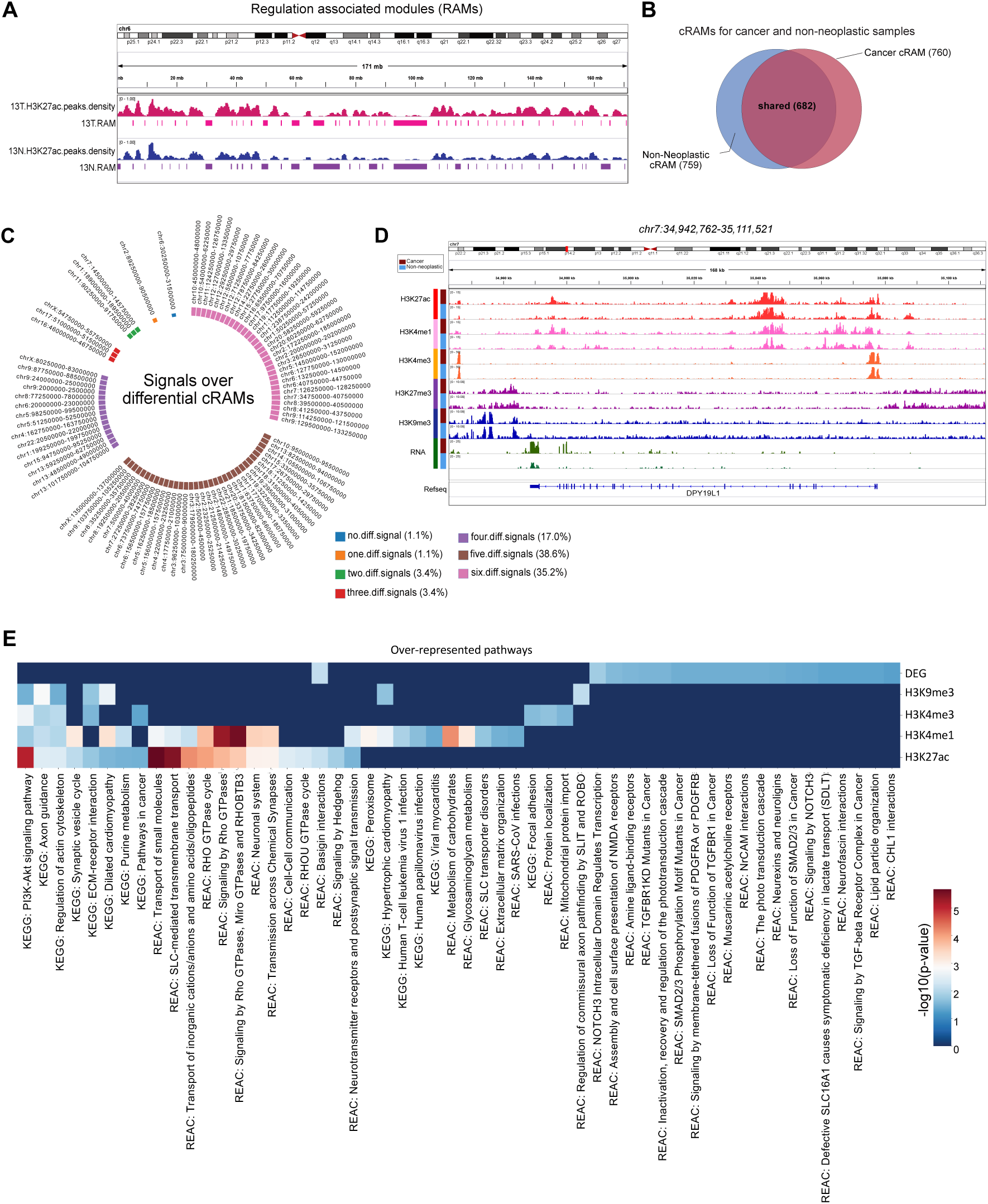
Differential regulatory associated modules (RAMs) analysis. **(A)** Genome browser view of individual RAMs on the chr6 in tumor and non-neoplastic samples. **(B)** Consensus RAMs (cRAMs) in tumor and non-neoplastic groups. **(C)** Differential epigenetic and expression signals in the tumor or normal tissue-specific cRAMs. **(D)** Genome browser view of multiple DMERs with differential cRAM regions in chr7. **(E)** Over-represented pathways in the cRAM regions overlapping epigenomic or transcriptomic signals.

### Gene regulatory network analysis reveals transcription factors and lncRNAs with altered roles

Finally, to elucidate the underlying regulators for the DEG patterns, we constructed the gene regulatory network (GRN) analyses for individual samples by integrating H3K27ac and RNA-seq data using the Taiji pipeline ^5^. Briefly, Taiji predicted transcription factor (TF) binding sites using known motifs in the active promoter and enhancer regions with H3K27ac marks, then linked enhancers to their putative interacting promoters as predicted by EpiTensor ^48^, then assembled all TF-gene pairs into a genetic network, and finally calculated the PageRank score that reflects the global importance for every node (TF or gene) in the network (see **Methods**).

By comparing PageRank scores for TFs between tumor and non-neoplastic groups, we identified 68 differentially-PageRanked TFs (|log2FC| >= 0.3, and *p*-value <= 0.05 by two-sided Mann-Whitney U test) (**Figure 5A** and **5B**), suggesting shifted global impacts of these TFs ^5,49^. Notably, 9 out of these TFs belonged to the Zinc-finger protein (ZNF) family, known to play crucial roles in tumorigenesis and progression ^50^. We successfully identified the previously verified ZFX ^51^ and ZNF322 ^52^, demonstrating the reliability of our approach. **Figure 5C** illustrated the over-represented pathways enriched in the significantly altered TFs. As expected, cancer-related pathways including “Transcriptional misregulation activities in cancers” in the KEGG database were identified. Additionally, multiple regulatory activities related to processes such as “SUMOylation” and “activation of HOX genes” were also involved.

**Figure 5.**
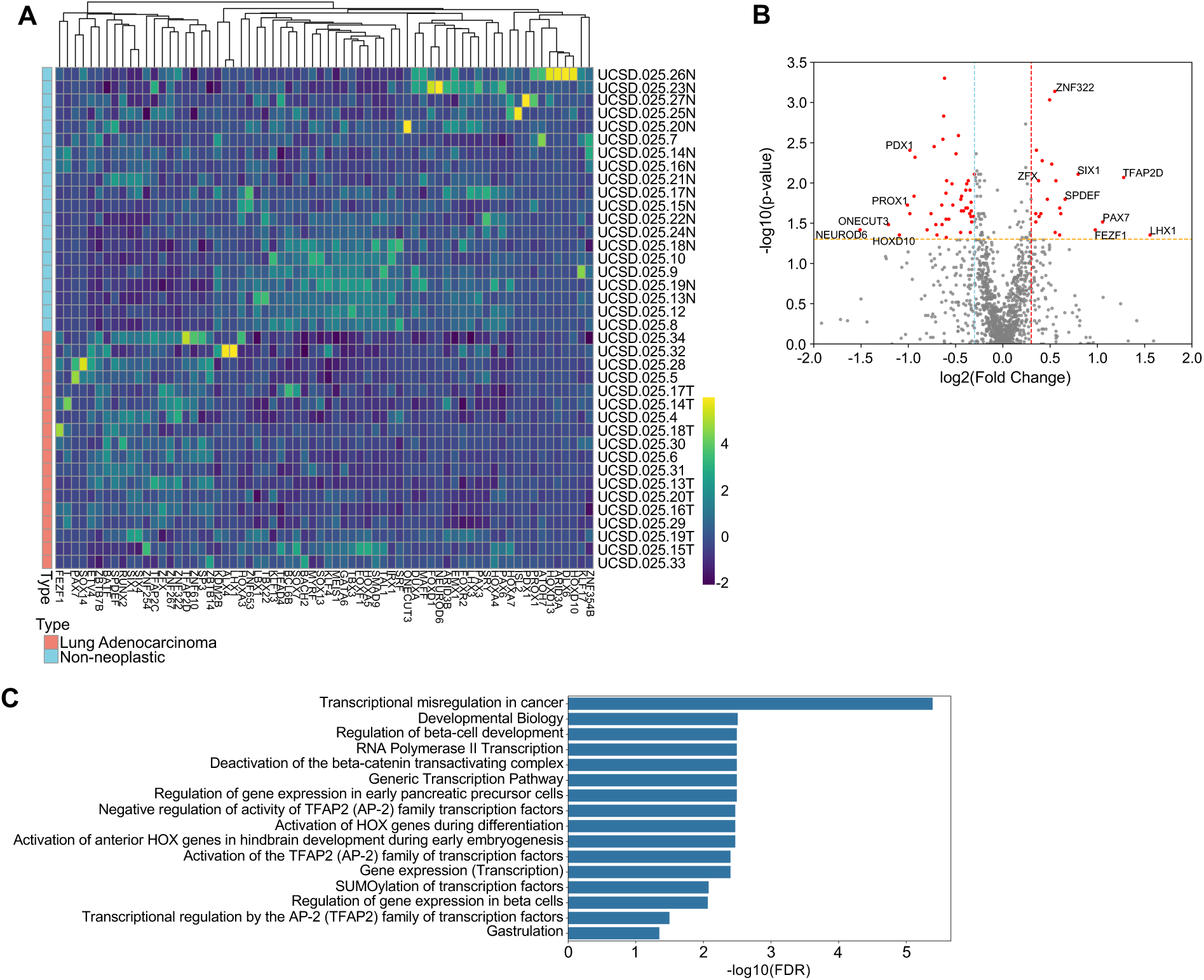
Gene regulatory network (GRN) analysis using Taiji. **(A)** Heatmap for the transcription factors (TFs) with significantly changed global importance (PageRank score). **(B)** Identification of the differentially PageRanked TFs. **(C)** Over-represented pathways for the differentially PageRanked TFs.

Among the altered transcription factors (TFs) identified by Taiji, SPDEF exhibited a high fold change (FC) of > 1.5 (**Figure 5B**). We examined the expression level of SPDEF by comparing its RPM in the tumor versus non-neoplastic groups and consistently, SPDEF displayed higher expression in the tumor samples (**Figure 6A**, *p*-value = 0.022 by Mann-Whitney U test). SPDEF was reported as a transcription factor exhibiting higher expression for tumors in brain, breast, and lung, compared with the corresponding normal tissues ^53,54^. We investigated SPDEF’s high-ranking regulatees (**Figure 6B**, see Methods) and also discovered its local gene regulatory network (**Figure 6C**). The Taiji regulatee analysis offers a comprehensive insight into various biomolecules (including genes, snRNAs, and lncRNAs) that play significant roles in lung cancer, even if their differential expression was not evident in RNA-seq experiments. The reliability of this method was underscored by the substantial literature support for more than 77% of SPDEF’s regulatees, highlighting their close interaction with NSCLC (**Figure 6B**). These regulatees encompass a diverse range of biomolecules, including early-stage biomarkers (ATXN10 ^55^), validated therapeutic targets (TCP1 ^56^), small-nuclear RNAs with prognostic potentials (SNORA7B ^57,58^), and genes that control migration and invasion of NSCLC (SERINC2 ^59^ and VANGL2 ^60^). For instance, the long noncoding RNA (lncRNA) RP11-523H20.3 ^61^ has been demonstrated to be related with the proliferation and metastasis of NSCLC. Conversely, other lncRNAs such as RP11-77P16.4 and RP11-927P21.4 represent novel candidates whose roles in lung cancer are yet to be fully studied (**Figure 6B**). RP11-773H22.4 is another promising novel participant in lung cancer, particularly notable due to its location within EpiSig Section 2 and the differential cRAM. Moreover, long intergenic non-coding RNAs (lincRNAs) like AC144450.1 can be identified by observing the integrative signals. It has increased H3K4me1 and H3K27ac signals and is located within cancer-specific differential cRAM (**Figure 6B**). Pathways that were associated with NSCLC development were found to be over-represented by SPDEF regulatees, highlighting their potential relevance in the disease progression. Specifically, “SUMOylation of ubiquitinylation proteins” pathway emerged due to the escalated activities of PIAS1 ^62^, SEH1L, and NUP37 ^63^; and “Regulation of IFN gamma signaling” pathway for PIAS1 ^62^, PTPN2 ^64^, and STAT1 ^65^ activities (**Figure 6D**).

**Figure 6.**
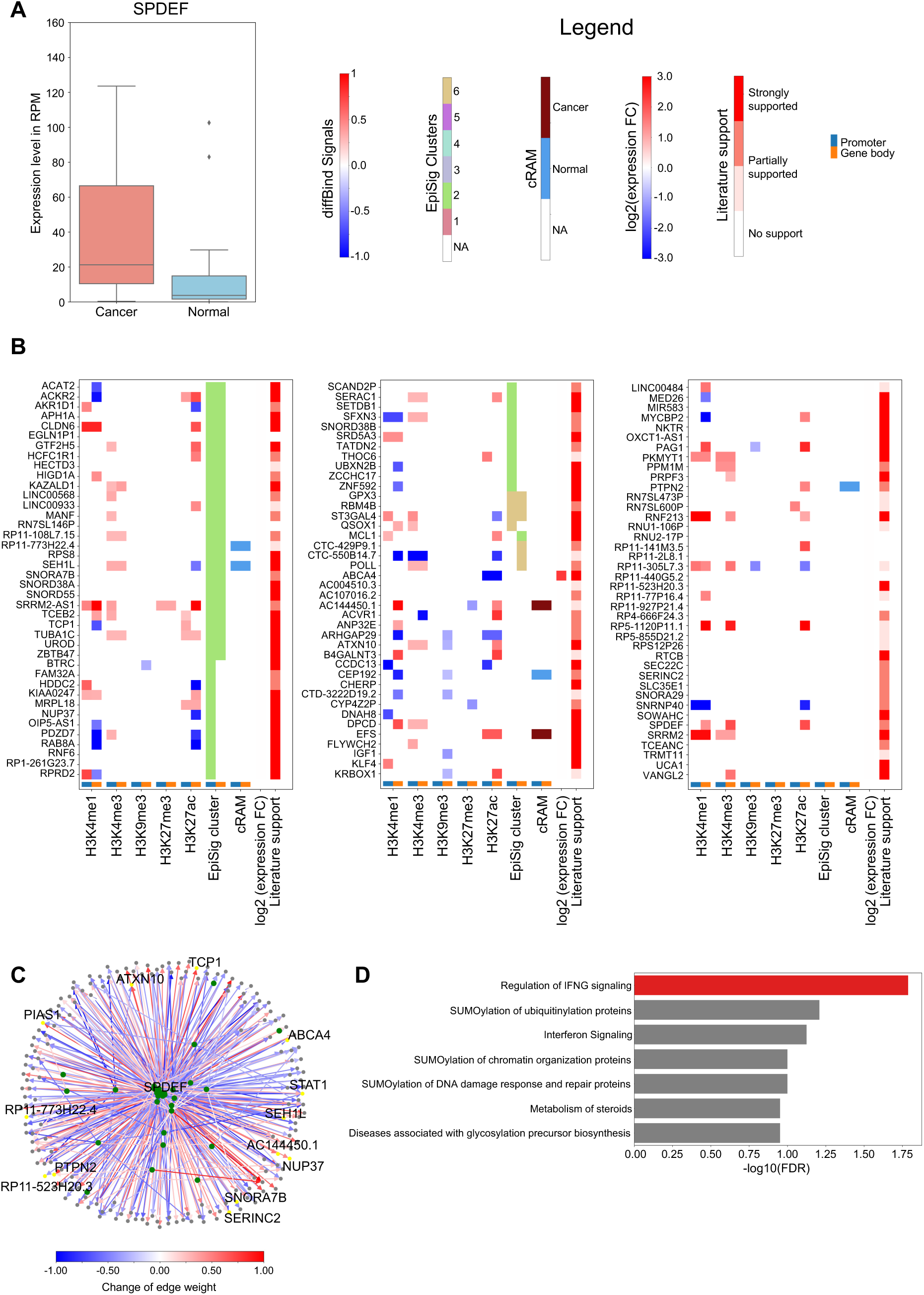
Regulatee analysis for SPDEF. **(A)** Boxplot of SPDEF’s expression levels in the tumor and non-neoplastic samples, *p* = 0.022 by Mann-Whitney U test. **(B)** Features and literature evidence for the complete list of SPDEF’s top regulatee genes. **(C)** Graph of gene regulatory activities between SPDEF and its target genes. Green nodes represent transcription factors, and blue nodes represent regulatees. Edge color indicates the difference of edge weights between tumor and non-neoplastic groups. (**D**) Over-represented pathways for SPDEF and its top 20% regulatees.

Further examples included SIX1 ^66^ and TFAP2D ^67^, along with their regulatees, demonstrating the diverse array of biomolecules identified by the Taiji analysis that were implicated in NSCLC development and progression (**Figure S6** and **S7).**

In summary, integrative analysis of significant TFs and their regulatees using Taiji facilitated the identification of a broad spectrum of biomolecules (such as early-stage biomarkers and well-studied therapeutic targets), along with genomic regions of marked with variable features, providing a comprehensive view of the regulatory landscape of NSCLC’s driving mechanisms. We derived a list of 1,414 biomolecules from all sources in our analyses that might be significant in the NSCLC (**Table S3**). This list is comprised of genes ranked in the following priority: (1) 96 DEGs (**Figure S8**), (2) 67 significant TFs (with one additional significant TF already included in the DEG list), (3) 238 regulatees controlled by multiple TFs, implying their central positions in the GRN, (4) 488 regulatees within EpiSig sections which contain multiple representative DMERs, (5) 38 regulatees within differential cRAM regions, implying that these regions had H3K27ac peaks that govern the organizations of chromatin activities, and (6) a total of other 487 strongly altered regulatees with no other obvious epigenomic signals.

In addition, we identified NSCLC-associated genetic regions as EpiSig regions harboring the above-mentioned genes. We counted the number of genes within these regions and extracted the top 5%. As a result, 5,639 genomic regions spanning 27 EpiSig clusters were found (**Table S4**).

## Discussion

In this work, the application of low-input technologies (MOWChIP-seq and Smart-seq2) enabled the simultaneous and high-quality profiling of five histone marks and transcriptome using a small quantity of tissue samples (∼ 20 mg). We established a robust and complete workflow for comprehensive multiomic characterization of tissue samples, offering a powerful tool for investigating the epigenomic dynamics associated with lung tumorigenesis.

Previous epigenomic studies on patient-derived tumor samples primarily focused on the open chromatin states ^8^ and active histone modifications ^7,12^, with limited exploration of repressive histone marks such as H3K27me3 and H3K9me3 due to technical challenges, although there is proved significance associated with these marks ^68^ and bivalent domains characterized by both active and repressive marks ^69^. By leveraging low-input technologies, our study expanded the scope of profiling of five histone marks, including three active marks and two repressive marks, alongside the transcriptome across all tumor and control samples.

Our comprehensive dataset enabled segregation of genome-wide co-modified epigenomic and transcriptomic regions into six distinct sections, including two repressed ones. While both H3K27me3 and H3K9me3 are repressive marks in nature, H3K9me3 is considered a permanent mark of repression, whereas H3K27me3 is more temporary^70^. Our integrative analysis revealed that H3K27me3 was enriched in Section IV, coupled with weak H3K4me3 and H3K4me1, making no significant contribution to cRAM alterations. In contrast, differentially modified H3K9me3 peaks were enriched in Section V, accounting for 31% of differential cRAMs, indicating that chromatin structure changes in NSCLC extend beyond the single epigenetic mark modifications. Overall, our results underscore that high-quality mapping of both active and repressive marks is important for generating a complete picture of the epigenetic landscape.

Our H3K27ac maps enabled further analysis on the organizations of chromosomal activities, by examining regulation associated modules (RAMs). In this work, we identified 88 differential cRAMs, effectively summing up the changes in RAM boundaries from tumorigenesis. Notably, 91% of differential cRAMs exhibited enrichment for multiple types of DMERs and DEGs, highlighting the relevance to the differences between tumor and non-neoplastic samples. Pathway analysis of sections within differential cRAMs revealed unique functional roles, including many cancer-related pathways, underscoring the utility of differential cRAMs in extracting informative genomic trends. Overall, our findings support the robustness of RAMs as modules of chromatin organization, with common patterns observed in cRAMs shared by tumor and non-neoplastic samples, while pathologically relevant changes are concentrated in differential cRAMs specific to each sample type. These results demonstrated the potential of RAMs as a valuable tool for analyzing and comparing epigenomic and transcriptomic datasets, providing insights into the underlying dynamics associated with tumorigenesis ^39^.

Integrating transcriptomic and chromatin organization data, we constructed a gene regulatory network (GRN) for each patient sample and computed the global importance of transcription factors (TFs) by calculating their PageRank scores, using the Taiji software. Differentially-PageRanked TFs were analyzed and their crucial functions in tumorigenesis were reported. Pathway analysis on these top TFs and their regulatees revealed common cancer-inducing pathways including “SUMOylation”, “activation of HOX genes”, and “interferon gamma signaling” pathways, echoing the findings based on differential cRAM analyses. The validation of top regulatees for example TFs (SPDEF, SIX1, and TFAP2D) further confirmed the reliability of our method, with over 70% of the selected genes supported by literature. Notably, the regulatees encompassed a diverse range of biomolecules, many of which showed no significant signals from expression or ChIP-seq experiments alone, highlighting the added value of our integrative approach.

The significance of our study lies in its contribution to understanding the complexity of solid tumors, which comprise cancerous and non-neoplastic cells with diverse roles in tumor development ^71^. Our data incorporate contributions from various cell types, enabling systems-level comparison and analyses with direct clinical relevance for patient stratification. Overall, our efforts in comprehensive multiomic profiling, particularly involving a large number of histone marks, pave the way for a more precise understanding of chromatin state dynamics during tumorigenesis, offering insights into potential therapeutic targets and strategies for cancer treatment.

## Author contributions

W.W. and C.L. conceived and supervised the project. W.W., C.L., Z.L., and L.Z. designed the experiments. Z.L. conducted MOWChIP-seq assays and Z.Z. conducted Smart-seq2 assays. P.W. and L.Z. conducted the data analysis. P.W. and Z.L. contributed to the data quality analysis. P.W., L.Z., Z.L., C.L., and W.W. wrote the manuscript. All authors proofread the manuscript and provided feedback.

## Data Availability

The raw ChIP-seq and RNA-seq data on the human lung tissues were deposited in dbGaP under accession number phs003113.v1.p1. The processed ChIP-seq and RNA-seq data can be accessed via Gene Expression Omnibus (GEO) under accession number GSE230932.

## Methods

### Human frozen lung tissues biopsies

The de-identified human frozen archived non-small cell lung cancer (lung adenocarcinoma) (N = 18) and non-neoplastic lung biopsies (N = 20) were obtained from Biorepository & Tissue Technology Shared Resource at Moores Cancer center, UC San Diego. UC San Diego and Virginia Tech IRB office reviewed our study proposal and determined the study status as “Not Human Subjects Research”. All the cancer samples were primary lung adenocarcinoma samples in pathological stage I and II (**Table S1**). The de-identified frozen samples were sealed in 1.8 mL cryovials and safely stored in liquid nitrogen vapor phase before epigenetic and transcriptomic profiling experiments.

### Isolation of nuclei

Samples were shipped to Virginia Tech on dry ice and stored in liquid nitrogen vapor phase until use. Thin slices (∼ 15 mg) were cut from the tissue on dry ice and transferred to a grinder (D9063, Sigma-Aldrich) pre-cooled on ice. The following steps were performed on ice and centrifugation performed at 4°C. 3 ml of ice-cold nuclei extraction buffer [0.32 M sucrose, 5 mM CaCl_2_, 3 mM Mg(Ac)_2_, 0.1 mM EDTA, 10 mM tris-HCl, and 0.1% Triton X-100, with 30 µl of PIC (P8340, Sigma-Aldrich), 3 µl of 100 mM PMSF, and 3 µl of 1 M dithiothreitol added before use] was added to the grinder. Tissue was homogenized by slowly douncing 20 times with pestle A and 30 times with pestle B. Crude nuclei was filtered through a 40 µm cell strainer into a 15 ml tube to remove debris. The mixture was centrifuged at 1000*g* for 10 min. Supernatant was removed and the crude nuclei pellet was resuspended in 500 µl nuclei extraction buffer and transferred to a 1.5 ml tube. 750 µl of 50% iodixanol, 7.5 µl of PIC, 0.75 µl of 100 mM PMSF and 0.75 µl of 1M dithiothreitol were added and mixed by gently pipetting and then inverting the tube three times. The mixture was centrifuged at 10,000*g* for 20 min and the supernatant was removed. The purified nuclei pellet was resuspended in 200 µl of Dulbecco’s phosphate-buffered saline (DPBS).

### MNase digestion of chromatin

Resuspended nuclei were counted and diluted to 4×10^6^/ml. 2 µl of PIC, 2 µl of 100 mM PMSF and 200 µl lysis buffer [4% Triton X-100, 100 mM tris-HCl, 100 mM NaCl, and 30 mM MgCl2] were added to 200 µl of nuclei suspension, mixed by vortexing and incubated at room temperature for 10 min. 20 µl of 100 mM CaCl_2_ and 5 µl of 100U MNase (88216, Thermo Fisher Scientific) were added, mixed by vortexing and incubated at room temperature for 10 min. 44 µl of 0.5 M EDTA was then added, mixed by vortexing and incubated on ice for 10 min. The mixture was centrifuged at 16,100*g* for 5 min at 4°C. Supernatant containing chromatin fragments (∼ 440 μl) were transferred to a new 1.5 ml tube and placed on ice. 30 µl of chromatin solution was used for each ChIP assay.

### Preparation of immunoprecipitation beads

50 µl of protein A Dynabeads (10001D, Invitrogen) were used in 10 MOWChIP assays for each tissue sample. Dynabeads were washed twice with 200 µl of IP buffer (20 mM tris-HCl, pH 8.0, 140 mM NaCl, 1 mM EDTA, 0.5 mM EGTA, 0.1% (w/v) sodium deoxycholate, 0.1% SDS, 1% (v/v) Triton X-100) and resuspended in 1.5 ml of IP buffer. They were then divided into 5 equal 300 µl portions in 1.5 ml tubes, each containing beads for 2 replicate assays. The following antibodies were added: anti-H3K27ac (ab4729, Abcam), 0.125 µg/assay; anti-H3K4me1 (39297, Active Motif), 0.5 µl/assay; anti-H3K4me3 (ab8580, Abcam), 0.125 µg/assay; anti-H3K27me3 (39155, Active Motif), 0.5 µg/assay; anti-H3K9me3 (ab8898, Abcam), 0.125 µg/assay. Beads were incubated overnight on a rotator at 4°C. Beads were washed with 200 µl of IP buffer twice prior to MOWChIP and resuspended in 10 µl of IP buffer for loading into the device.

### MOWChIP

Fabrication of the MOWChIP device, control system setup and operation of the device were done following our previously published protocol ^18^.

### Purification of ChIP DNA

After MOWChIP, IP beads with bound ChIP DNA were collected from the device, washed with 200 µl of IP buffer and resuspended in 200 µl of DNA elution buffer (10 mM tris-HCl, 50 mM NaCl, 10 mM EDTA, and 0.03% SDS). 2 µl of 20 mg/ml proteinase K was added and incubated at 65°C for 1 h. ChIP DNA was extracted and purified by phenol-chloroform extraction and ethanol precipitation. Air-dried DNA pellet was dissolved in 8 µl of low EDTA TE buffer. Input DNA was prepared by digesting 6 µl of chromatin solution directly using proteinase K without ChIP process, using the same extraction and purification protocol.

### ChIP Library preparation, quantification and sequencing

Libraries were prepared using Accel NGS 2S Plus DNA library kit (IDT) following the manufacturer’s protocol with the following modification: 5% EvaGreen dye (Biotium) was added to the amplification mixture to monitor progress. Amplification was stopped once a 3000 RFU increase from baseline was observed. Library was eluted to 10 µl of low EDTA TE buffer and stored at -20°C. ChIP enrichment of the libraries was quantified using qPCR (with primers in **Table S5**). Library concentration was quantified using Kapa Library Quantification kit (Roche). Libraries were pooled for sequencing by Illumina HiSeq 4000 in SR75 mode.

### RNA-seq

Tissue slices (∼ 5 mg) were placed in 200 µl of RNAlater-ICE frozen tissue transition solution (AM7030, Thermo Fisher Scientific) prechilled to -80°C. Tissue was transitioned at -20°C overnight. Each RNA-seq library was prepared from 10ng RNA extracted from the tissue. Total RNA was extracted using RNeasy Mini Kit (74104, Qiagen) and RNase-Free DNase Set (79254, Qiagen). Reverse transcription reaction was performed following the Smart-seq2 ^72^ protocol with minor modifications. Extracted total RNA was resuspended in 4.5 μL of RNase-free water with 5% RNase inhibitor (40 U/μL) added. 2 μL of oligo-dT primer (10 μM), 2 μL of dNTP mix (10 mM) and 4.3 μL of total RNA solution were mixed, incubated at 72°C for 3 minutes and immediately put on ice. 1 μL of SuperScript II reverse transcriptase (200 U/μL), 0.5μL of RNase inhibitor (40 U/μl), 4μL of Superscript II first-strand buffer, 1 μL of DTT (100 mM), 4 μL of 5 M Betaine, 0.12 μL of 1 M MgCl_2_, 0.2 μL of TSO (100 μM), 0.88 μL of nuclease-free water were added. The reverse transcription (RT) reaction mix was then incubated at 42°C for 90 minutes, 10 cycles of 50°C for 2 minutes, 42°C for 2 minutes, and 70°C for 15 minutes. 25 μL KAPA HiFi HotStart ReadyMix, 0.5 μL IS PCR primers (10 μM), 2.5 μL Evagreen dye, and 2 μL nuclease-free water were added. The mixture was amplified with the following program: 98°C for 1 minute, 9–11 cycles of 98°C for 15 seconds, 67°C for 30 seconds and 72°C for 6 minutes. The cDNA was purified using 30 μL of SPRIselect beads and eluted in 5 μL of low EDTA TE buffer.

The RNA-seq library was made using tagmentation. 3 μL of home-made Tn5 was mixed with 3 μL of 50 μM pre-annealed P5/P7 transposon and incubated at 37°C for 1 hour.

Assembled P5 and P7 transposons were then mixed well to form Tn5 mix. 1 μL cDNA (600 pg) and 1 μL of tagmentation buffer (TNP92110, Lucigen) were added to 8 μL Tn5 mix, and the mixture was incubated at 37°C for 1 hour. 1 μL of 10X stop buffer (TNP92110, Lucigen) was then added to the mixture to quench tagmentation. 11 μL SPRIselect beads were used to purify tagmented cDNA. The cDNA was then eluted in 9.5 μL of low EDTA TE buffer. 25 μL of KAPA HiFi HotStart ReadyMix was incubated at 98°C for 30 seconds, mixed with 9.5 μL tagmented cDNA and incubated at 72°C for 5 minutes. 1.5 μL of 100 μM P5 primer, 1.5 μL of 100 μM P7 primer, 10 μL nuclease-free water and 2.5 μL Evagreen were then added and amplified with the following program: 98°C for 30 seconds; 10-12 cycles of 98°C for 10 seconds, 63°C for 30 seconds, and 72°C for 30 seconds. Amplified cDNA libraries were purified with 50 μL SPRIselect beads and eluted in 8 μL of low EDTA TE buffer. Library concentration was quantified using Kapa Library Quantification kit (Roche). RNA-seq libraries were sequenced with Illumina HiSeq 4000 platform.

## Data Analysis

### RNA-seq data processing

The raw fastq files were trimmed using Trim Galore ^73^, and then aligned against the hg19 reference genome using the STAR software ^74^. The mean number of uniquely mapped reads was 11,462,974 for RNA-seq libraries (**Table S2**). Reads with Q30 were retained for further analysis. Quality assessment reports for each RNA-seq library were generated using QualiMap software ^75^. Gene expression for each sample was quantified using the featureCounts software ^76^.

Differentially expressed genes (DEGs) between the tumor and non-neoplastic samples were identified using the DESeq2 package ^23^ in R. A typical DEG was defined based on a two-fold change in gene expression levels and an adjusted *p*-value of <= 0.05. For further gene expression analysis, we applied the regularized log transformation on the read counts using the DESeq2 package.

### ChIP-seq data processing

Raw ChIP-seq data for H3K4me1, H3K4me3, H3K9me3, H3K27ac, and H3K27me3 were aligned to the hg19 genome using bwa (0.7.17-r1188) ^77^. After the deduplication step, the uniquely aligned reads with MAPQ >= 10 were retained for further processing.

The mean numbers of uniquely mapping reads were: 12,191,347 for H3K27ac; 12,151,681 for H3K4me1; 16,998,758 for H3K4me3; 25,536,891 for H3K27me3; and 37,784,615 for H3K9me3. Narrow peaks (H3K27ac, H3K4me1, and H3K4me3) were identified using MACS2 (2.2.7.1) with a *q*-value of <= 0.05 ^78^. Broad peaks (H3K9me3 and H3K27me3) were called using Homer ^79^ findPeaks in histone mode with a *p*-value of <= 0.05. On average, for each sample, there were 90,164 peaks for H3K27ac, 130,793 peaks for H3K4me1, 38,855 peaks for H3K4me3, 32,444 peaks for H3K27me3, and 54,511 peaks for H3K9me3 (**Table S2**).

Library enrichment and complexity were assessed through quality control evaluation, including the fraction of reads that fell into peak regions (FRiP). Replicated peaks were identified by merging peaks from the two replicates for each sample, where replicated peaks were defined as those mutually overlapped by at least 50%.

Differentially binding sites for individual histone marks between tumor and non-neoplastic samples were identified using the DiffBind package in R with default parameters. A typical differentially binding site was identified with a false discovery rate (FDR) <= 0.05.

### Genome-wide multi-omics clustering analysis by EpiSig

The genome-wide epigenetic signals and gene expressions were clustered using an unsupervised learning method EpiSig ^34,35^. To ensure comparability across all sequencing data from tumor and non-neoplastic samples, normalization was performed by adjusting the total sequencing depth. Regions in the ENCODE blacklist ^80^ were excluded from the processed data.

The ChIP-seq signals subtracting inputs, and the RNA-seq data were input into the EpiSig pipeline. The EpiSig algorithm segmented the genome into 5-kb bins and detected enriched signals within these bins in the whole genome across histone modifications and gene expressions among all the samples. The 5-kb bins with similar signal patterns were clustered into EpiSig clusters. To better interpret the EpiSig cluster outputs, individual EpiSig clusters were further summarized into large sections using the K-Means clustering approach, with the optimal value of *K* determined by elbow plotting.

### Identification of Epi-DMERs

“Epi-DMERs” were identified as DMER loci with differential signals (H3K27ac, H3K4me1, H3K4me3, H3K27me3, H3K9me3, and RNA-seq) residing in any of the EpiSig clusters.

### Genome-wide annotation of the EpiSig regions

We downloaded CpG, TSSs, 1 kb upstream of TSS, gene exons, intronic regions, intergenic regions, 3’ UTR, and 5’ UTR coordinate files for the hg19 genome from the UCSC genome browser database ^81^. Overlaps between these regions and the regions in the EpiSig clusters were examined.

### Identification of regulatory associated modules (RAMs) in individual samples

H3K27ac narrow-peak density was calculated using a sliding window with a step size of 250 kb and flanking size of 500 kb for each window in every sample. RAM boundaries (valley/minima on the smoothing curves) and peaks (summit/maxima on the smoothing curves) were detected using the “findRAM” tool ^39^.

### Identification of consensus RAMs (cRAMs)

We first identified RAMs in 20 non-neoplastic samples with replicates or 18 tumor samples with replicates, using a step size of 250 kb. We then counted the percentage of genomic regions identified as RAM boundaries in tumor or non-neoplastic groups.

Regions with percentage >= 25% were considered as consensus RAM (cRAM) boundaries in each group respectively. Since a cRAM was required to span at least 500 kb, cRAM boundaries located < 250 kb apart from each other were merged.

### Identification of globally important transcription factors (TFs) using Taiji

For both tumor and non-neoplastic samples, processed H3K27ac narrow peaks and raw RNA-seq read counts were used as inputs for the Taiji software. Promoter-enhancer interactions were predicted as the top 10% of all contacts identified from EpiTensor ^48^, an unsupervised learning algorithm. Putative TF binding motifs were obtained from the CIS-BP database ^82^. The Taiji pipeline (v1.3) was executed with default parameters ^5^. For each transcription factor (TF), We performed Mann-Whitney U test by comparing its PageRank score between tumor and non-neoplastic groups to derive a *p*-value.

Subsequently, 68 significantly important TFs were identified with |log2FC| >= 0.3 and *p*- value <= 0.05 in this way (**Figure 5B**, **Table S3**).

### Identification of significantly altered target genes (regulatees)

Edges (representing TF -> regulatee) in the gene regulatory networks were extracted from the Taiji output for each sample. Weak and unreliable edges were filtered out if their weight ranks fell below 50%, and the percentile ranks of the remaining edges were calculated. Edges that were not consistently detected in at least five individual samples were considered unreliable and thus discarded. For each remaining edge, we conducted Mann-Whitney U test by comparing its percentile rank scores between the tumor and non-neoplastic group to derive a *p*-value. The edge weight difference was calculated as the mean percentile rank score of the tumor group subtracted from that of the non-neoplastic group. Edges with *p*-value <= 0.05 were selected, and their corresponding end nodes were identified as the significantly altered regulatees. The comprehensive list of all significant regulatees was provided in **Table S3**.

### Comprehensive listing of all promising NSCLC-associated biomolecules

For genes (biomolecules) identified through any of the methods below: (1) DEGs, (2) significant TFs, and (3) significant regulatees (top 200 for each TF, respectively), the following information was annotated: (1) overlap with all 5 ChIP-seq signals (H3K27ac, H3K4me1, H3K4me3, H3K27me3, H3K9me3), (2) overlap with EpiSig clusters, (3) overlap with differential cRAMs, and (4) literature support indicating significance in lung cancer.

All overlapping steps were achieved with the bedtools ^83^ software, considering either the gene body or the promoter regions (up to 1 kb upstream of the TSS). Literature support was categorized into four levels: 0 for no reported evidence, 1 for weaker support indicating general physiological functions, 2 for stronger support implying importance in the lung cancer; and 4 for solid support for its functions or relationship with the NSCLC.

Genes were ranked according to the following priorities: (1) 96 DEGs, (2) 67 significant TFs, (3) 238 regulatees controlled by multiple TFs, implying their central positions in the GRN, (4) 488 regulatees within EpiSig sections which contain multiple representative DMERs, (5) 38 regulatees within differential cRAM regions, implying that these regions had H3K27ac peaks that govern the organizations of chromatin activities, and (6) a total of other 487 strongly altered regulatees (**Table S3**).

NSCLC-associated genetic regions were identified as EpiSig regions harboring the above-mentioned genes. The top 5% of these EpiSig regions were selected (**Table S4**).

## References

1. Flavahan, W. A., Gaskell, E. & Bernstein, B. E. Epigenetic plasticity and the hallmarks of cancer. Science 357, (2017).

2. Waddington, C. H. The Strategy of the Genes (Vol. George Allen & Unwin): London. (1957).

3. Suvà, M. L., Riggi, N. & Bernstein, B. E. Epigenetic Reprogramming in Cancer. Science 339, 1567–1570 (2013).

4. Feinberg, A. P., Koldobskiy, M. A. & Göndör, A. Epigenetic modulators, modifiers and mediators in cancer aetiology and progression. Nat. Rev. Genet. 17, 284–299 (2016).

5. Zhang, K., Wang, M., Zhao, Y. & Wang, W. Taiji: System-level identification of key transcription factors reveals transcriptional waves in mouse embryonic development. Sci Adv 5, eaav3262 (2019).

6. Mirhadi, S. et al. Integrative analysis of non-small cell lung cancer patient-derived xenografts identifies distinct proteotypes associated with patient outcomes. Nat. Commun. 13, 1811 (2022).

7. Pomerantz, M. M. et al. Prostate cancer reactivates developmental epigenomic programs during metastatic progression. Nat. Genet. 52, 790–799 (2020).

8. Corces, M. R. et al. The chromatin accessibility landscape of primary human cancers. Science 362, (2018).

9. Belinsky, S. A. Gene-promoter hypermethylation as a biomarker in lung cancer. Nat. Rev. Cancer 4, 707–717 (2004).

10. Seligson, D. B. et al. Global levels of histone modifications predict prognosis in different cancers. Am. J. Pathol. 174, 1619–1628 (2009).

11. Barlési, F. et al. Global histone modifications predict prognosis of resected non small-cell lung cancer. J. Clin. Oncol. 25, 4358–4364 (2007).

12. Li, Q.-L. et al. Genome-wide profiling in colorectal cancer identifies PHF19 and TBC1D16 as oncogenic super enhancers. Nat. Commun. 12, 6407 (2021).

13. Kelley, D. Z. et al. Integrated Analysis of Whole-Genome ChIP-Seq and RNA-Seq Data of Primary Head and Neck Tumor Samples Associates HPV Integration Sites with Open Chromatin Marks. Cancer Res. 77, 6538–6550 (2017).

14. Cohen, A. J. et al. Hotspots of aberrant enhancer activity punctuate the colorectal cancer epigenome. Nat. Commun. 8, 14400 (2017).

15. Lomberk, G. et al. Distinct epigenetic landscapes underlie the pathobiology of pancreatic cancer subtypes. Nat. Commun. 9, 1978 (2018).

16. Della Chiara, G. et al. Epigenomic landscape of human colorectal cancer unveils an aberrant core of pan-cancer enhancers orchestrated by YAP/TAZ. Nat. Commun. 12, 2340 (2021).

17. Cao, Z., Chen, C., He, B., Tan, K. & Lu, C. A microfluidic device for epigenomic profiling using 100 cells. Nat. Methods 12, 959–962 (2015).

18. Zhu, B. et al. MOWChIP-seq for low-input and multiplexed profiling of genome-wide histone modifications. Nat. Protoc. 14, 3366–3394 (2019).

19. Liu, Z., et al. nMOWChIP-seq: low-input genome-wide mapping of non-histone targets. NAR Genom Bioinform 4, lqac030 (2022).

20. Picelli, S. et al. Smart-seq2 for sensitive full-length transcriptome profiling in single cells. Nat. Methods 10, 1096–1098 (2013).

21. Marinov, G. K., Kundaje, A., Park, P. J. & Wold, B. J. Large-scale quality analysis of published ChIP-seq data. G3 4, 209–223 (2014).

22. Ross-Innes, C. S. et al. Differential oestrogen receptor binding is associated with clinical outcome in breast cancer. Nature 481, 389–393 (2012).

23. Love, M. I., Huber, W. & Anders, S. Moderated estimation of fold change and dispersion for RNA-seq data with DESeq2. Genome Biol. 15, 550 (2014).

24. Raudvere, U. et al. g:Profiler: a web server for functional enrichment analysis and conversions of gene lists (2019 update). Nucleic Acids Res. 47, W191–W198 (2019).

25. Mehlen, P., Delloye-Bourgeois, C. & Chédotal, A. Novel roles for Slits and netrins: axon guidance cues as anticancer targets? Nat. Rev. Cancer 11, 188–197 (2011).

26. Biankin, A. V. et al. Pancreatic cancer genomes reveal aberrations in axon guidance pathway genes. Nature 491, 399–405 (2012).

27. Wang, W.-J. et al. Dolutegravir derivative inhibits proliferation and induces apoptosis of non-small cell lung cancer cells via calcium signaling pathway. Pharmacol. Res. 161, 105129 (2020).

28. Zhang, Y.-L., Wang, R.-C., Cheng, K., Ring, B. Z. & Su, L. Roles of Rap1 signaling in tumor cell migration and invasion. Cancer Biol Med 14, 90–99 (2017).

29. Boettner, B. & Van Aelst, L. Control of cell adhesion dynamics by Rap1 signaling. Curr. Opin. Cell Biol. 21, 684–693 (2009).

30. Vasan, N., Boyer, J. L. & Herbst, R. S. A RAS Renaissance: Emerging Targeted Therapies for KRAS-Mutated Non–Small Cell Lung Cancer. Clin. Cancer Res. 20, 3921–3930 (2014).

31. Papadimitrakopoulou, V. Development of PI3K/AKT/mTOR Pathway Inhibitors and Their Application in Personalized Therapy for Non–Small-Cell Lung Cancer. J. Thorac. Oncol. 7, 1315–1326 (2012).

32. Fang, J. Y. & Richardson, B. C. The MAPK signalling pathways and colorectal cancer. Lancet Oncol. 6, 322–327 (2005).

33. Eke, I. & Cordes, N. Focal adhesion signaling and therapy resistance in cancer. Semin. Cancer Biol. 31, 65–75 (2015).

34. Ai, R. et al. Comprehensive epigenetic landscape of rheumatoid arthritis fibroblast-like synoviocytes. Nat. Commun. 9, 1921 (2018).

35. Hon, G., Ren, B. & Wang, W. ChromaSig: a probabilistic approach to finding common chromatin signatures in the human genome. PLoS Comput. Biol. 4, e1000201 (2008).

36. Kim, T. et al. Comparative characterization of 3D chromatin organization in triple-negative breast cancers. Exp. Mol. Med. 54, 585–600 (2022).

37. Vilarrasa-Blasi, R. et al. Dynamics of genome architecture and chromatin function during human B cell differentiation and neoplastic transformation. Nat. Commun. 12, 651 (2021).

38. Brock, M. V., Herman, J. G. & Baylin, S. B. Cancer as a manifestation of aberrant chromatin structure. Cancer J. 13, 3–8 (2007).

39. Zheng, L. & Wang, W. Regulation associated modules reflect 3D genome modularity associated with chromatin activity. Nat. Commun. 13, 5281 (2022).

40. Kanehisa, M., Furumichi, M., Tanabe, M., Sato, Y. & Morishima, K. KEGG: new perspectives on genomes, pathways, diseases and drugs. Nucleic Acids Res. 45, D353– D361 (2017).

41. Gillespie, M. et al. The reactome pathway knowledgebase 2022. Nucleic Acids Res. 50, D687–D692 (2022).

42. Naba, A., Clauser, K. R., Lamar, J. M., Carr, S. A. & Hynes, R. O. Extracellular matrix signatures of human mammary carcinoma identify novel metastasis promoters. Elife 3, e01308 (2014).

43. Naba, A. et al. Characterization of the Extracellular Matrix of Normal and Diseased Tissues Using Proteomics. J. Proteome Res. 16, 3083–3091 (2017).

44. Hebert, J. D. et al. Proteomic Profiling of the ECM of Xenograft Breast Cancer Metastases in Different Organs Reveals Distinct Metastatic Niches. Cancer Res. 80, 1475–1485 (2020).

45. Parker, A. L. et al. Extracellular matrix profiles determine risk and prognosis of the squamous cell carcinoma subtype of non-small cell lung carcinoma. Genome Med. 14, 126 (2022).

46. Jeon, H.-S. & Jen, J. TGF-^2^ Signaling and the Role of Inhibitory Smads in Non-small Cell Lung Cancer. J. Thorac. Oncol. 5, 417–419 (2010).

47. Park, S. et al. Crizotinib attenuates cancer metastasis by inhibiting TGFβ signaling in non-small cell lung cancer cells. Exp. Mol. Med. 54, 1225–1235 (2022).

48. Zhu, Y. et al. Constructing 3D interaction maps from 1D epigenomes. Nat. Commun. 7, 10812 (2016).

49. Yu, B. et al. Erratum: Epigenetic landscapes reveal transcription factors that regulate CD8+ T cell differentiation. Nat. Immunol. 18, 705 (2017).

50. Cassandri, M. et al. Zinc-finger proteins in health and disease. Cell Death Discov 3, 17071 (2017).

51. Jen, J. & Wang, Y.-C. Zinc finger proteins in cancer progression. J. Biomed. Sci. 23, 53 (2016).

52. Jen, J. et al. Oncogenic zinc finger protein ZNF322A promotes stem cell-like properties in lung cancer through transcriptional suppression of c-Myc expression. Cell Death Differ. 26, 1283–1298 (2019).

53. Song, J. et al. A Radioresponse-Related lncRNA Biomarker Signature for Risk Classification and Prognosis Prediction in Non-Small-Cell Lung Cancer. J. Oncol. 2021, 4338838 (2021).

54. Bao, K.-C. & Wang, F.-F. The role of SPDEF in cancer: promoter or suppressor. Neoplasma 69, 1270–1276 (2022).

55. Yin, L.-G. et al. Analysis of tissue-specific differentially methylated genes with differential gene expression in non-small cell lung cancers. Mol. Biol. 48, 694–700 (2014).

56. Carr, A. C. et al. Targeting chaperonin containing TCP1 (CCT) as a molecular therapeutic for small cell lung cancer. Oncotarget 8, 110273–110288 (2017).

57. Zhuo, Y. et al. Targeting SNORA38B attenuates tumorigenesis and sensitizes immune checkpoint blockade in non-small cell lung cancer by remodeling the tumor microenvironment via regulation of GAB2/AKT/mTOR signaling pathway. J Immunother Cancer 10, (2022).

58. Feng, S. et al. Identification of Six Novel Prognostic Gene Signatures as Potential Biomarkers in Small Cell Lung Cancer. Comb. Chem. High Throughput Screen. 26, 938– 949 (2023).

59. Zeng, Y. et al. SERINC2-knockdown inhibits proliferation, migration and invasion in lung adenocarcinoma. Oncol. Lett. 16, 5916–5922 (2018).

60. Xu, Z. et al. Circ-IGF1R inhibits cell invasion and migration in non-small cell lung cancer. Thorac. Cancer 11, 875–887 (2020).

61. Li, X. et al. Genetic Variants of CLPP and M1AP Are Associated With Risk of Non-Small Cell Lung Cancer. Front. Oncol. 11, 709829 (2021).

62. Constanzo, J. D. et al. PIAS1-FAK Interaction Promotes the Survival and Progression of Non-Small Cell Lung Cancer. Neoplasia 18, 282–293 (2016).

63. Huang, L. et al. NUP37 silencing induces inhibition of cell proliferation, G1 phase cell cycle arrest and apoptosis in non-small cell lung cancer cells. Pathol. Res. Pract. **216**, 152836 (2020).

64. Wang, C.-C. et al. Novel Potential Therapeutic Targets of PTPN Families for Lung Cancer. J Pers Med 12, (2022).

65. Li, J., Yu, B., Song, L., Eschrich, S. & Haura, E. B. Effects of IFN-gamma and Stat1 on gene expression, growth, and survival in non-small cell lung cancer cells. J. Interferon Cytokine Res. 27, 209–220 (2007).

66. Huang, S. et al. SIX1 Predicts Poor Prognosis and Facilitates the Progression of Non-small Lung Cancer via Activating the Notch Signaling Pathway. J. Cancer 13, 527–540 (2022).

67. Yao, R., Zhou, L., Guo, Z., Zhang, D. & Zhang, T. Integrative molecular analyses of an individual transcription factor-based genomic model for lung cancer prognosis. Dis. Markers 2021, 5125643 (2021).

68. Brykczynska, U. et al. Repressive and active histone methylation mark distinct promoters in human and mouse spermatozoa. Nat. Struct. Mol. Biol. 17, 679–687 (2010).

69. Bernstein, B. E. et al. A bivalent chromatin structure marks key developmental genes in embryonic stem cells. Cell 125, 315–326 (2006).

70. Kim, J. & Kim, H. Recruitment and biological consequences of histone modification of H3K27me3 and H3K9me3. ILAR J. 53, 232–239 (2012).

71. Egeblad, M., Nakasone, E. S. & Werb, Z. Tumors as organs: complex tissues that interface with the entire organism. Dev. Cell 18, 884–901 (2010).

72. Picelli, S. et al. Full-length RNA-seq from single cells using Smart-seq2. Nat. Protoc. 9, 171–181 (2014).

73. Krueger, F. TrimGalore: A Wrapper around Cutadapt and FastQC to Consistently Apply Adapter and Quality Trimming to FastQ Files, with Extra Functionality for RRBS Data. (Github).

74. Dobin, A. et al. STAR: ultrafast universal RNA-seq aligner. Bioinformatics 29, 15–21 (2013).

75. García-Alcalde, F. et al. Qualimap: evaluating next-generation sequencing alignment data. Bioinformatics 28, 2678–2679 (2012).

76. Liao, Y., Smyth, G. K. & Shi, W. featureCounts: an efficient general purpose program for assigning sequence reads to genomic features. Bioinformatics 30, 923–930 (2014).

77. Li, H. & Durbin, R. Fast and accurate short read alignment with Burrows-Wheeler transform. Bioinformatics 25, 1754–1760 (2009).

78. Zhang, Y. et al. Model-based analysis of ChIP-Seq (MACS). Genome Biol. 9, R137 (2008).

79. Heinz, S. et al. Simple combinations of lineage-determining transcription factors prime cis-regulatory elements required for macrophage and B cell identities. Mol. Cell 38, 576–589 (2010).

80. Amemiya, H. M., Kundaje, A. & Boyle, A. P. The ENCODE Blacklist: Identification of Problematic Regions of the Genome. Sci. Rep. 9, 1–5 (2019).

81. Navarro Gonzalez, J., et al. The UCSC Genome Browser database: 2021 update. Nucleic Acids Res. 49, D1046–D1057 (2020).

82. Weirauch, M. T. et al. Determination and inference of eukaryotic transcription factor sequence specificity. Cell 158, 1431–1443 (2014).

83. Quinlan, A. R. & Hall, I. M. BEDTools: a flexible suite of utilities for comparing genomic features. Bioinformatics 26, 841–842 (2010).

