## Supplementary Information for "Comprehensive multimodal and multiomic profiling reveals epigenetic and transcriptional reprogramming in lung tumors"

### Supplementary Figures

#### ChIP-seq

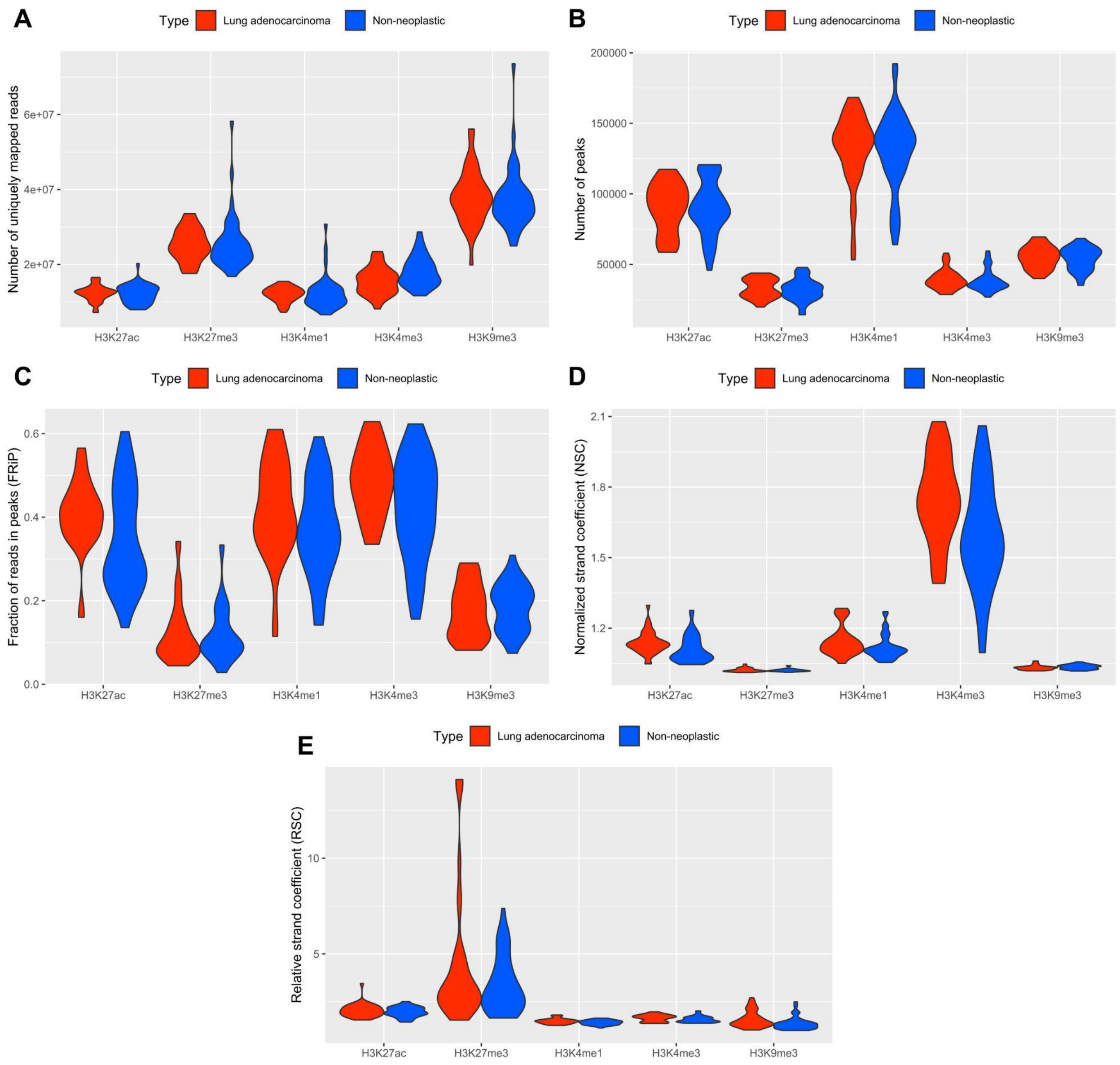


**Figure S1. Five quality control metrics for ChIP-seq data of H3K27ac, H3K27me3, H3K4me1, H3K4me3 and H3K9me3 from the lung tumor and non-neoplastic samples. (A)** Number of uniquely mapped reads. **(B)** Number of the peaks. **(C)** Fraction of reads in peaks (FRiP). **(D)** Normalized strand coefficient (NSC). **(E)** Relative strand coefficient (RSC).

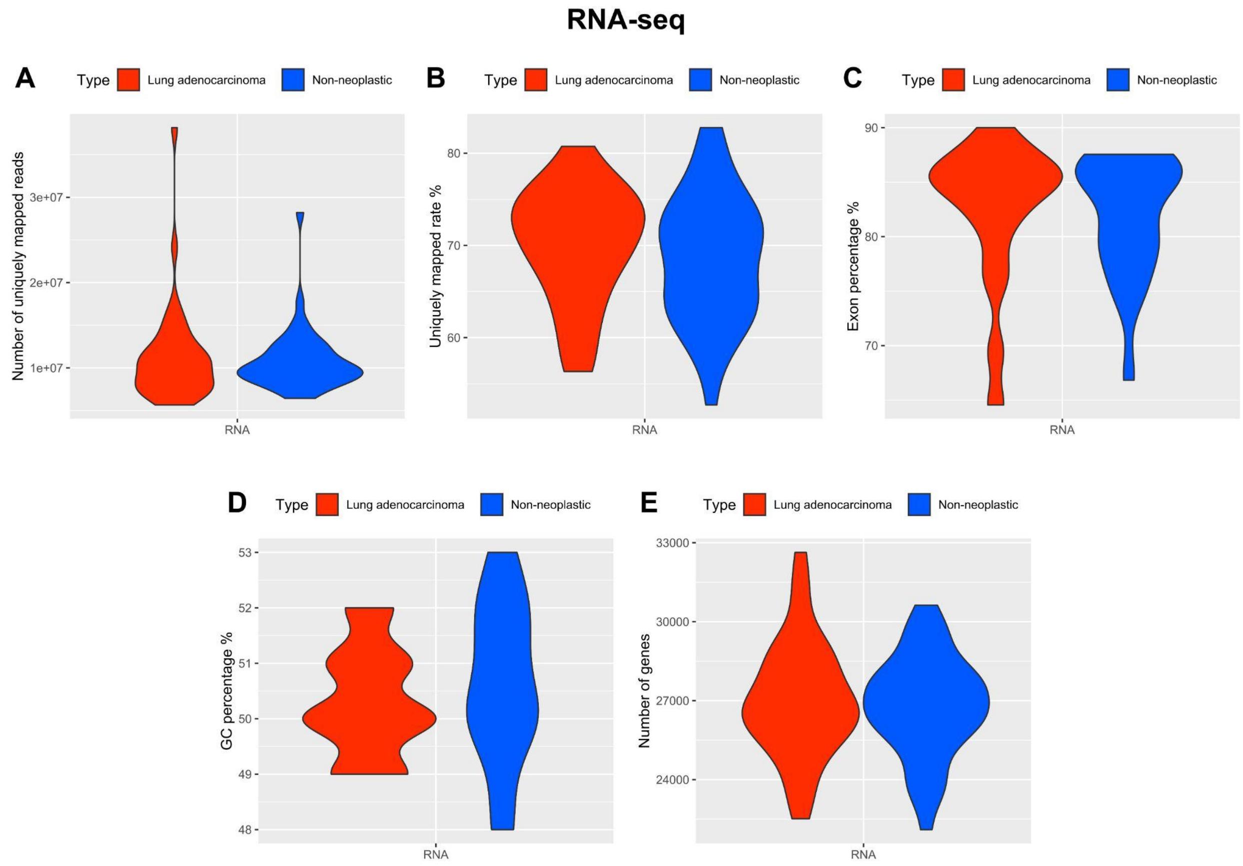


**Figure S2. Five quality control metrics for RNA-seq data from the lung tumor and non-neoplastic samples. (A)** Number of uniquely mapped reads. **(B)** Uniquely mapping rate. **(C)** Percentage of reads overlapping exons. **(D)** GC content. **(E)** Number of genes.

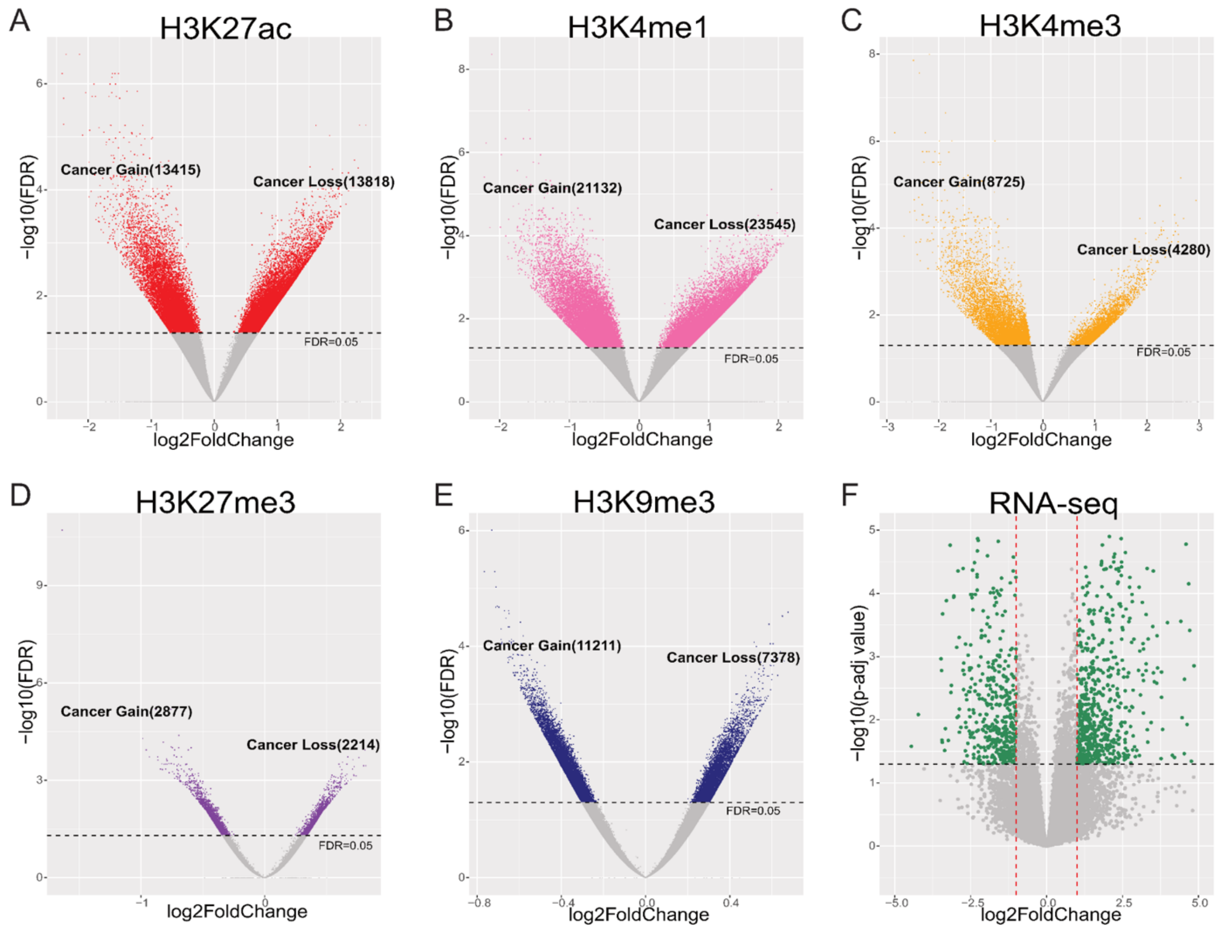


**Figure S3. Identification of differentially modified epigenetic regions (DMERs) and differentially expressed genes (DEGs) between tumor and non-neoplastic samples. (A)** H3K27ac DMERs. **(B)** H3K4me1 DMERs. **(C)** H3K4me3 DMERs. **(D)** H3K27me3 DMERs. **(E)** H3K9me3 DMERs. **(F)** DEGs.

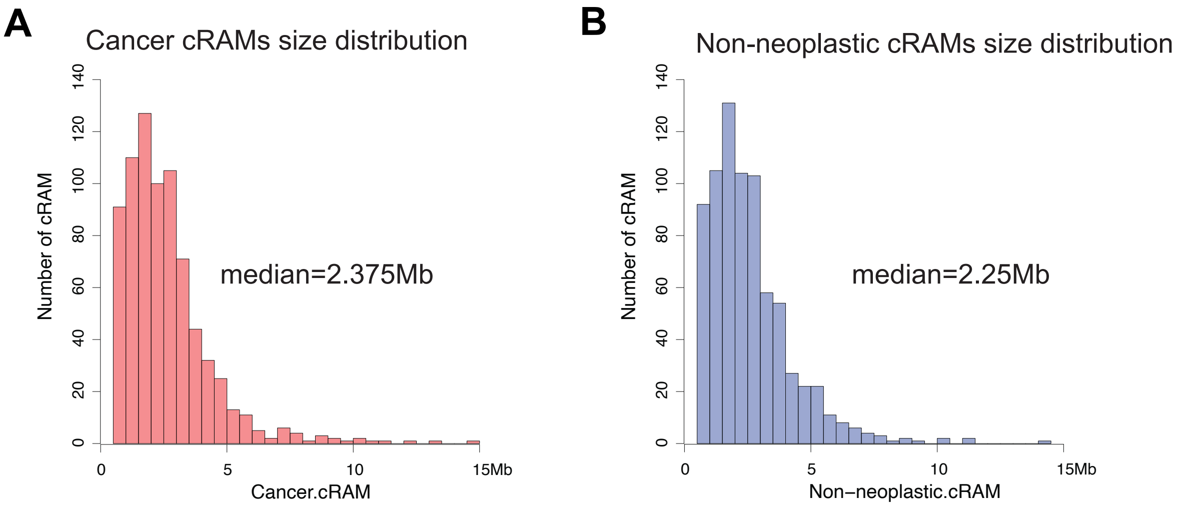


**Figure S4. Size distribution of cRAMs. (A)** tumor group, and **(B)** non-neoplastic group.

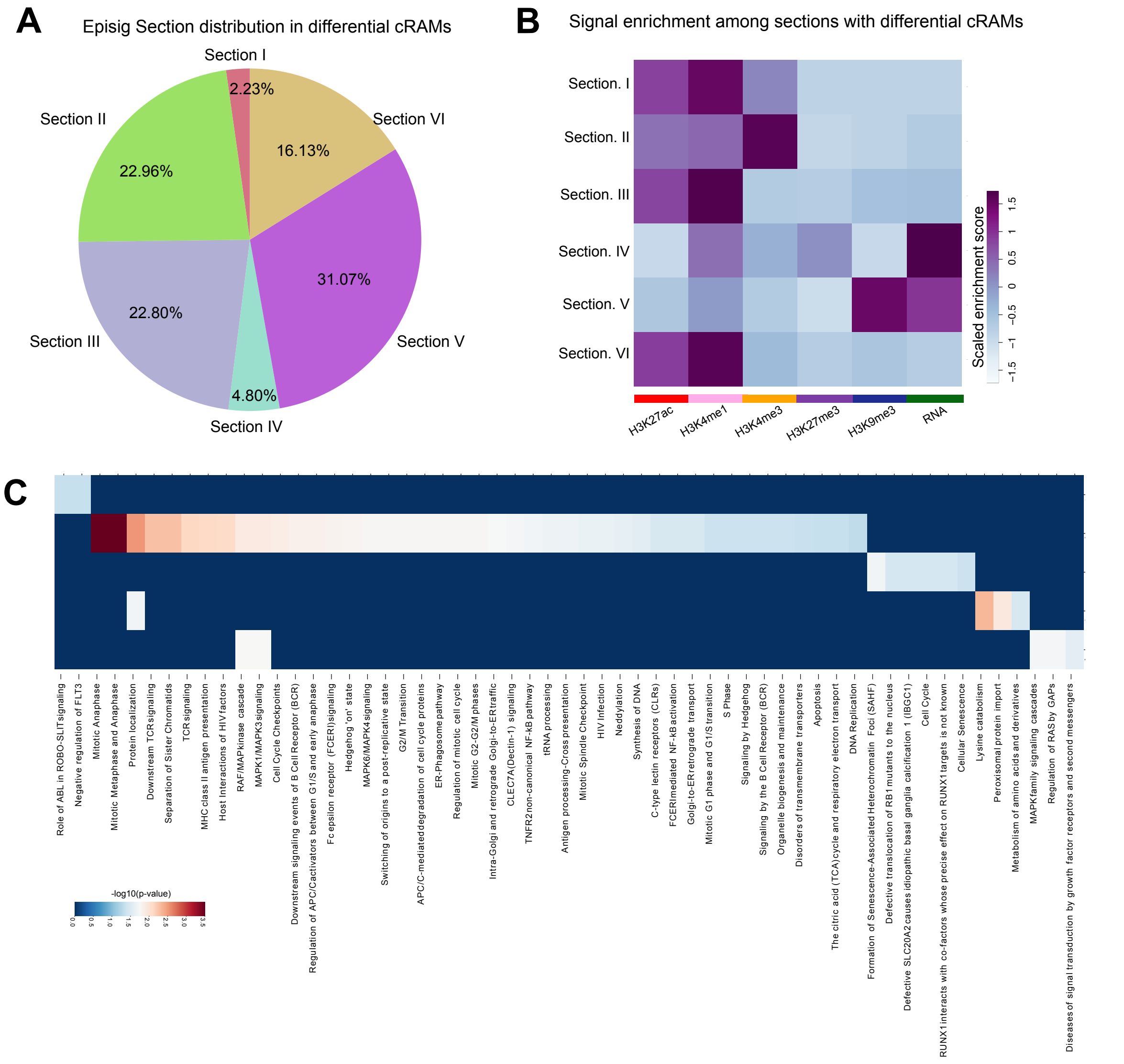


**Figure S5. Differential cRAMs in the EpiSig sections. (A)** Distribution of EpiSig sections in the differential cRAMs. **(B)** Enrichment of DMERs among the EpiSig sections with differential cRAMs. **(C)** Over-represented pathways in the EpiSig sections with differential cRAMs. Briefly, NSCLC samples' Section I showed enrichment with "Negative regulation of FLT3", a well-known biomarker in hematopoiesis studies; Section III with "defective translocation of RB1 mutants to the nucleus"; and Section VI with "MAPK family signaling cascade". Most importantly, Section II showed enrichment with critical cancer-related pathways such as "TCR signaling", "MAPK signaling pathways", "Apoptosis", "DNA replication", and "cell cycle checkpoints". Section V was mainly H3K9me3 with few annotated genes, showing no enrichment in the Reactome database.

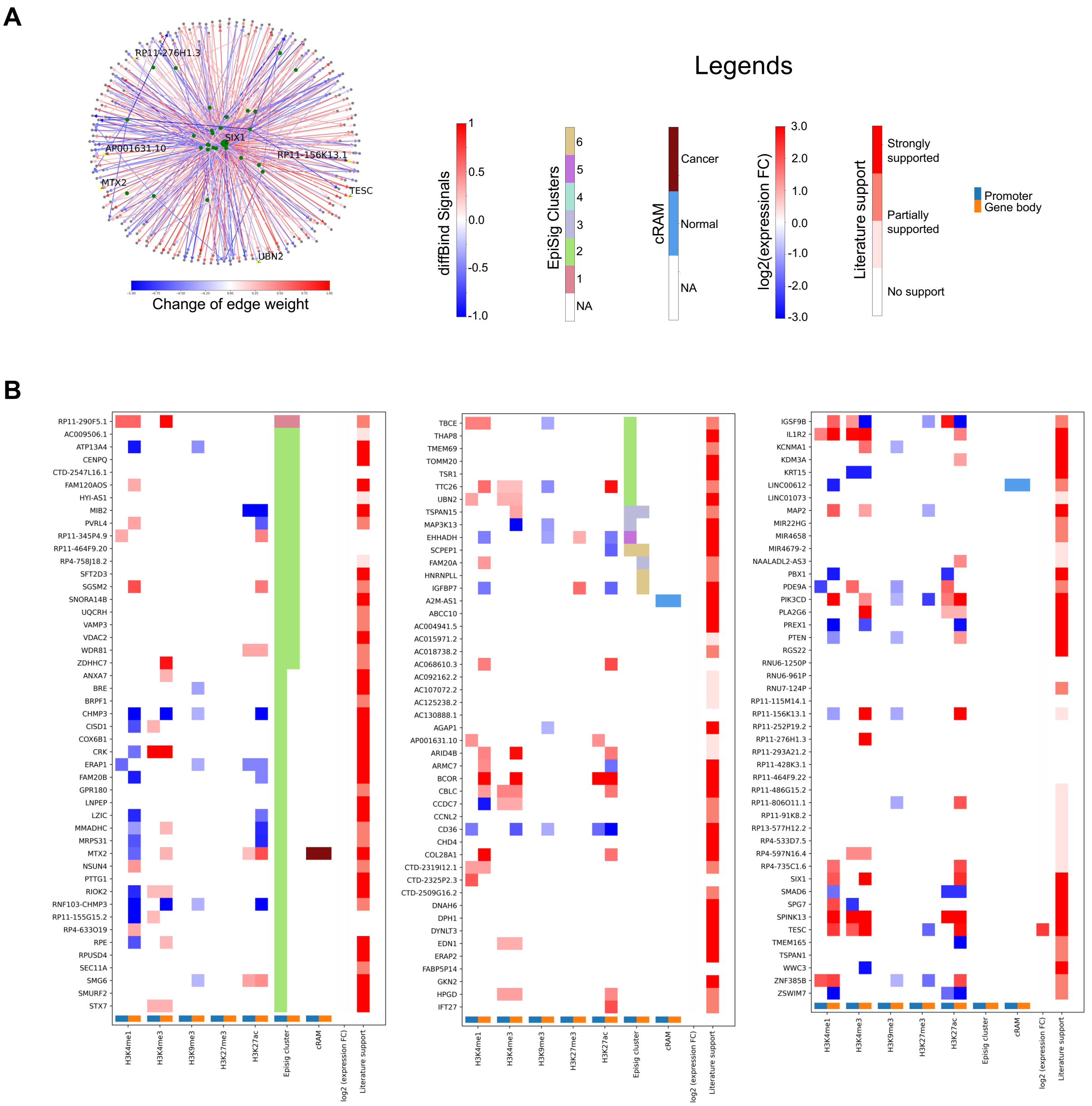


**Figure S6. Regulatee analysis for SIX1. (A)** Graph of gene regulatory activities between SIX1 and its target genes. Green nodes represent transcription factors, and blue nodes represent regulatees. Edge color indicates the difference of edge weights between cancer and normal sample groups. **(B)** Features and literature evidence for the complete list of SIX1's top regulatee genes.

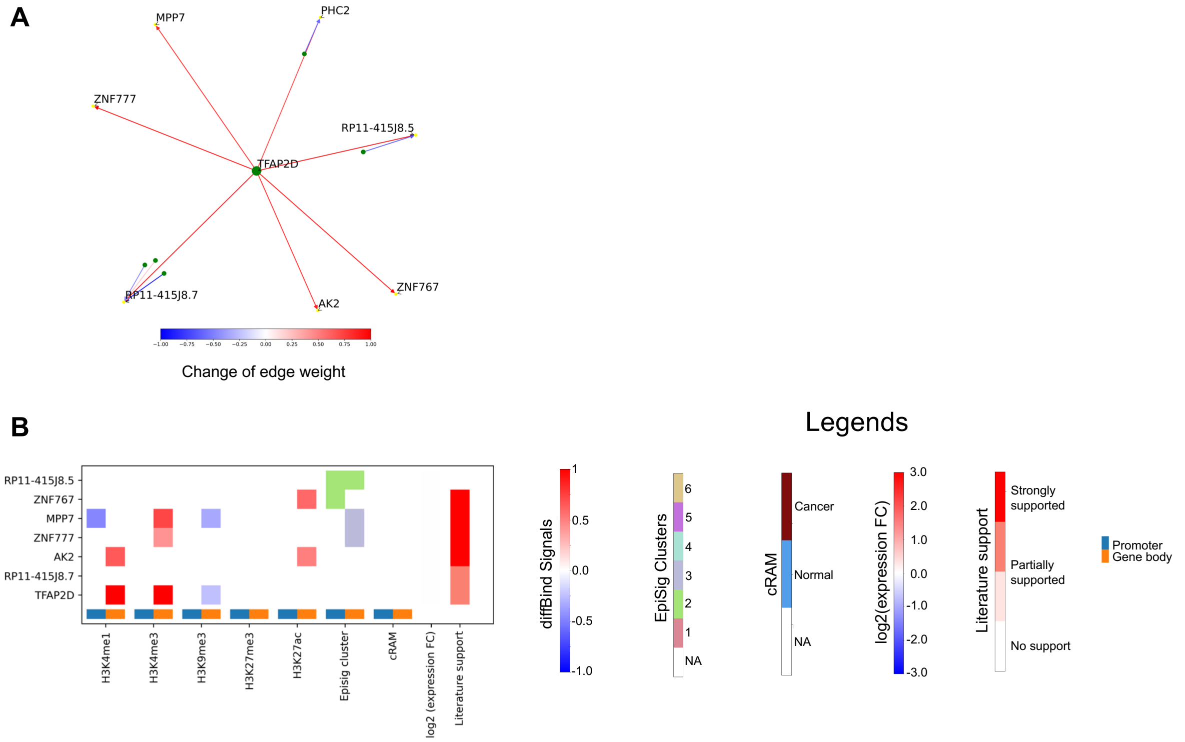


**Figure S7. Regulatee analysis for TFAP2D. (A)** Graph of gene regulatory activities between TFAP2D and its target genes. Green nodes represent transcription factors, and yellow nodes represent the emphasized regulates. Edge color indicates the difference of edge weights between cancer and normal sample groups. **(B)** Features and literature evidence for the complete list of TFAP2D's top regulatee genes.

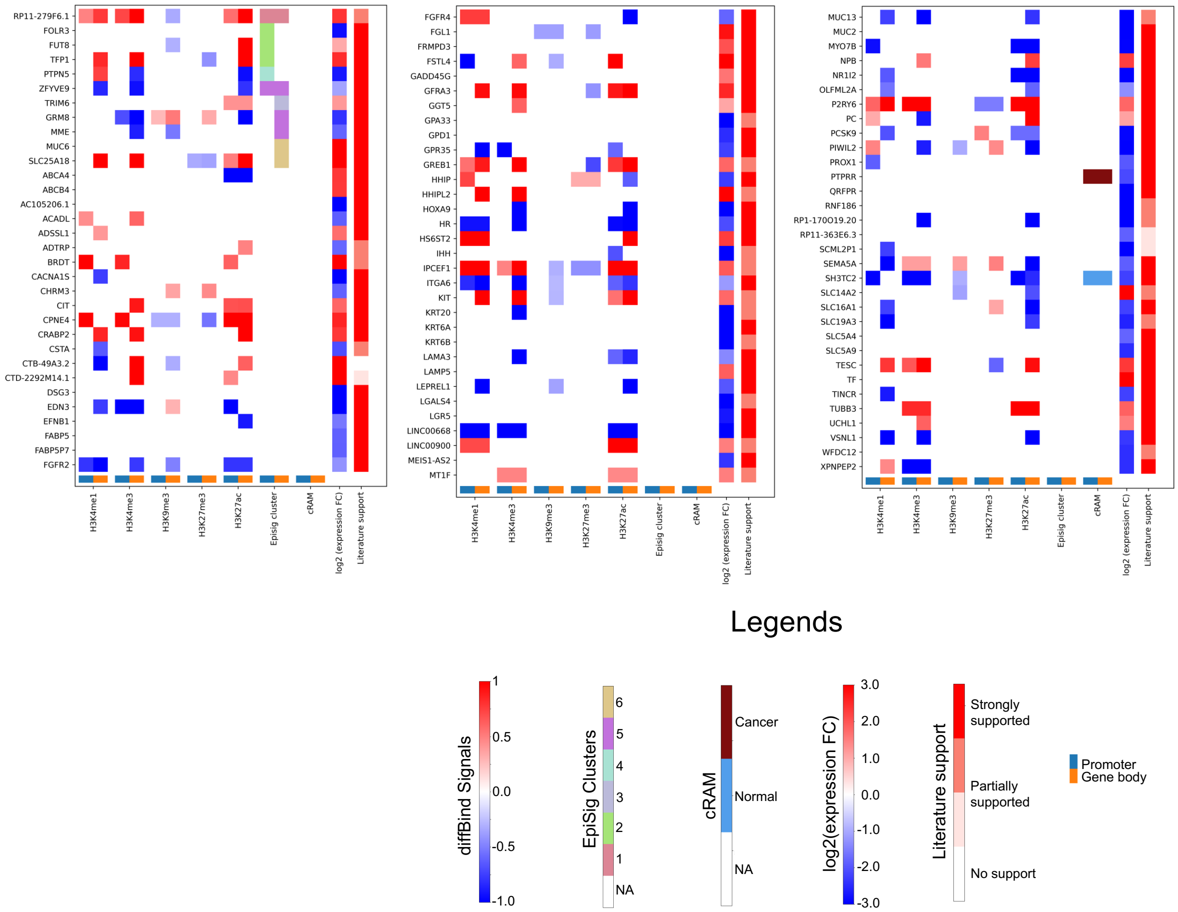


**Figure S8. Features and literature evidence for all DEGs.**

### Supplementary Tables

**Table S1. Tumor and non-neoplastic samples from patient lungs.** The numbers after "UCSD-025" differentiate various patients. Patients 13-20 contributed both tumor (T) and non-neoplastic (N) samples.

**Table S2. Metadata of ChIP-seq and RNA-seq data.**

**Table S3. List of 1,414 biomolecules important to NSCLC.**

**Table S4. List of 5,639 loci epigenetically important to NSCLC.**

**Table S5. Primers used in ChIP-qPCR for examining enrichment of the ChIP-seq libraries.**
